# PML-Dependent Memory of Type I Interferon Treatment Results in a Restricted Form of HSV Latency

**DOI:** 10.1101/2021.02.03.429616

**Authors:** Jon B. Suzich, Sean R. Cuddy, Hiam Baidas, Sara Dochnal, Eugene Ke, Austin R. Schinlever, Aleksandra Babnis, Chris Boutell, Anna R. Cliffe

**Author notes:** Correspondence to Anna R. Cliffe.

## Abstract

Herpes simplex virus (HSV) establishes latent infection in long-lived neurons. During initial infection, neurons are exposed to multiple inflammatory cytokines but the effects of immune signaling on the nature of HSV latency is unknown. We show that initial infection of primary murine neurons in the presence of type I interferon (IFN) results in a form of latency that is restricted for reactivation. We also found that the subnuclear condensates, promyelocytic leukemia-nuclear bodies (PML-NBs), are absent from primary sympathetic and sensory neurons but form with type I IFN treatment and persist even when IFN signaling resolves. HSV-1 genomes colocalized with PML-NBs throughout a latent infection of neurons only when type I IFN was present during initial infection. Depletion of PML prior to or following infection did not impact the establishment latency; however, it did rescue the ability of HSV to reactivate from IFN-treated neurons. This study demonstrates that viral genomes possess a memory of the IFN response during *de novo* infection, which results in differential subnuclear positioning and ultimately restricts the ability of genomes to reactivate.

## Introduction

Herpes simplex virus-1 (HSV-1) is a ubiquitous pathogen that persists in the form of a lifelong latent infection in the human host. HSV-1 can undergo a productive lytic infection in a variety of cell types; however, latency is restricted to post-mitotic neurons, most commonly in sensory, sympathetic and parasympathetic ganglia of the peripheral nervous system (Bloom, 2016). During latent infection, the viral genome exists as an episome in the neuronal nucleus, and there is considerable evidence that on the population level viral lytic gene promoters assemble into repressive heterochromatin (Cliffe et al., 2009, Knipe and Cliffe, 2008). The only region of the HSV genome that undergoes active transcription, at least in a fraction of latently infected cells, is the latency associated transcript (LAT) locus (Kramer and Coen, 1995, Stevens et al., 1987). Successful establishment of a latent gene expression program requires a number of molecular events, likely influenced by both cellular and viral factors, and is not uniform (Efstathiou and Preston, 2005). Significant heterogeneity exists in expression patterns of both lytic and latent transcripts in latently-infected neurons, as well as in the ability of latent genomes to reactivate in response to different stimuli (Proenca et al., 2008, Sawtell, 1997, Bertke et al., 2011, Nicoll et al., 2016, Ma et al., 2014, Catez et al., 2012, Maroui et al., 2016). This heterogeneity could arise from viral genome copy number, exposure to different inflammatory environments or intrinsic differences in the neurons themselves. Furthermore, there is growing evidence that heterogeneity in latency may ultimately be reflected in the association of viral genomes with different nuclear domains or cellular proteins (Catez et al., 2012, Maroui et al., 2016). However, what determines the subnuclear distribution of latent viral genomes is not known. In addition, it is currently unclear whether viral genome association with certain nuclear domains or cellular proteins results in an increased or decreased ability of the virus to undergo reactivation. The aim of this study was to determine whether the presence of interferon during initial HSV-1 infection can intersect with the latent viral genome to regulate the type of gene silencing and ultimately the ability to undergo reactivation. Because the fate of viral genomes and their ability to undergo reactivation can be readily tracked, latent HSV-1 infection of neurons also serves as an excellent system to explore how exposure to innate immune cytokines can have a lasting impact on peripheral neurons.

Latent HSV-1 genomes have been shown to associate with Promyelocytic leukemia nuclear bodies (PML-NBs) in mouse models of infection, as well as in human autopsy material (Catez et al., 2012, Maroui et al., 2016). PML-NBs are heterogenous, phase-separated nuclear condensates that have been associated with the transcriptional activation of cellular genes (Lallemand-Breitenbach and de The, 2010, Bernardi and Pandolfi, 2007, McFarlane et al., 2019, Wang et al., 2004, Kim and Ahn, 2015), but also can recruit repressor proteins, including ATRX, Daxx and Sp100, that promote transcriptional repression and inhibition of both DNA and RNA virus replication (Zhong et al., 2000, Garrick et al., 2004, Xu and Roizman, 2017, Everett and Chelbi-Alix, 2007, Bishop et al., 2006). In the context of lytic infection of non-neuronal cells, PML-NBs have been shown to closely associate with HSV-1 genomes (Maul et al., 1996, Maul, 1998), and the HSV-1 viral regulatory protein ICP0 is known to disrupt the integrity of these structures by targeting PML and other PML-NB associated proteins for degradation (Everett and Maul, 1994, Boutell et al., 2002, Chelbi-Alix and de The, 1999). PML-NBs entrapment of HSV-1 genomes during lytic infection of fibroblasts (Alandijany et al., 2018) is hypothesized to create a transcriptionally repressive environment for viral gene expression, as PML directly contributes to the cellular repression of ICP0-null mutant viruses (Everett et al., 2006). In the context of latency, neurons containing PML-encased latent genomes exhibit decreased expression levels of the LAT (Catez et al., 2012), suggesting that they are more transcriptionally silent than latent genomes localized to other nuclear domains and raising the question as to whether PML-NB-associated genomes are capable of undergoing reactivation. Studies have shown that replication-defective HSV genomes associated with PML-NBs are capable of derepressing following induced expression of ICP0 in fibroblasts (Cohen et al., 2018) and following addition of the histone deacetylase inhibitor trichostatin A (TSA) in cultured adult TG neurons (Maroui et al., 2016). However, it is not known if replication-competent viral genomes associated with PML-NBs are capable of undergoing reactivation triggered by activation of cellular signaling pathways in the absence of viral protein.

PML-NBs can undergo significant changes in number, size and localization depending on cell type, differentiation stage and cell-cycle phase, as well as in response to cellular stress and soluble factors (Lallemand-Breitenbach and de The, 2010, Bernardi and Pandolfi, 2007). Interferon (IFN) treatment directly induces the transcription of PML, Daxx, Sp100 and other PML-NB constituents, which leads to elevated protein synthesis and a robust increase in both size and number of PML-NBs (Chelbi-Alix et al., 1995, Stadler et al., 1995, Greger et al., 2005, Shalginskikh et al., 2013, Grotzinger et al., 1996). During HSV-1 infection, type I IFNs are among the first immune effectors produced and restrict HSV viral replication and spread both *in vitro* and *in vivo* through multiple pathways (Jones et al., 2003, Mikloska et al., 1998, Hendricks et al., 1991, Mikloska and Cunningham, 2001, Sainz and Halford, 2002). Although type I IFNs are elevated within peripheral ganglia during HSV-1 infection (Carr et al., 1998) and have been linked with control of lytic HSV-1 replication, whether type I IFN exposure modulates entry into latency is not known. Importantly, exposure to IFN and other cytokines has also been shown to generate innate immune memory or ‘trained immunity’ in fibroblasts and immune cells (Kamada et al., 2018, Moorlag et al., 2018), and PML-NBs themselves are potentially important in the host innate immune response. A previous study found that the histone chaperone HIRA is re-localized to PML-NBs in response to the innate immune defenses induced by HSV-1 infection, and in this context, PML was required for the recruitment of HIRA to ISG promoters for efficient transcription (McFarlane et al., 2019). Prior exposure to type I interferons has also been shown to promote a transcriptional memory response in fibroblasts and macrophages (Kamada et al., 2018). This interferon memory lead to faster and more robust transcription of ISGs following restimulation and coincided with acquisition of certain chromatin marks and accelerated recruitment of transcription and chromatin factors (Kamada et al., 2018). Thus far, long term memory of cytokine exposure has only been investigated in non-neuronal cells, but it is conceivable that neurons, being non-mitotic and long-lived cells, also possess unique long-term responses to prior cytokine exposure.

Although *in vivo* models are incredibly powerful tools to investigate the contribution of the host immune response to HSV infection, they are problematic for investigating how individual components of the host’s immune response specifically regulate neuronal latency. Conversely, *in vitro* systems provide a simplified model that lack many aspects of the host immune response. Therefore, to investigate the role of type I IFN on HSV-1 latency and reactivation, we utilized a model of latency in primary murine sympathetic neurons (Cliffe et al., 2015), which allowed us to manipulate conditions during initial HSV-1 infection and trigger synchronous robust reactivation. Using this model, we show that primary neurons isolated from mouse peripheral ganglia are largely devoid of PML-NBs but PML-NBs form following type I IFN exposure and persist even when ISG gene expression and production of other antiviral proteins have returned to baseline. Latency can be established in the presence of acyclovir, indicating that neither exogenous type I IFN nor PML-NBs are essential for HSV gene silencing and entry into latency in this model system. Importantly, the presence of IFNα specifically at the time of initial infection results in the entrapment of viral genomes in PML-NBs and a more restrictive form of latency that is less able to undergo reactivation. This study therefore demonstrates how the viral latent genome has a long-term memory of the innate response during *de novo* HSV infection that results in entrapment of genomes in PML-NBs and a more repressive form of latency.

## Results

### Interferon induces the formation of PML-NBs in primary sympathetic and sensory neurons isolated from postnatal and adult mice

We initially set out to investigate the contribution of PML-NBs to HSV latency and reactivation using primary sympathetic and sensory neurons that have been well characterized as *in vitro* models of HSV latency and reactivation (Camarena, 2011, Cliffe et al., 2015, Ives and Bertke, 2017, Wilcox and Johnson, 1987, Wilcox et al., 1990, Cuddy et al., 2020). In addition, primary neuronal systems allow for much more experimental control of specific conditions during *de novo* infection and can be easily manipulated either immediately prior to or following infection. Peripheral neurons were isolated from the superior cervical ganglia (SCG) or trigeminal ganglia (TG) from young (post-natal day; P1) or adult (>P28) mice and cultured for 6 days prior to staining. PML-NBs were defined as punctate nuclear structures by staining for PML protein. Strikingly, we observed that both SCG and TG neurons were largely devoid of PML-NBs (Fig. 1A).

**Fig. 1.**
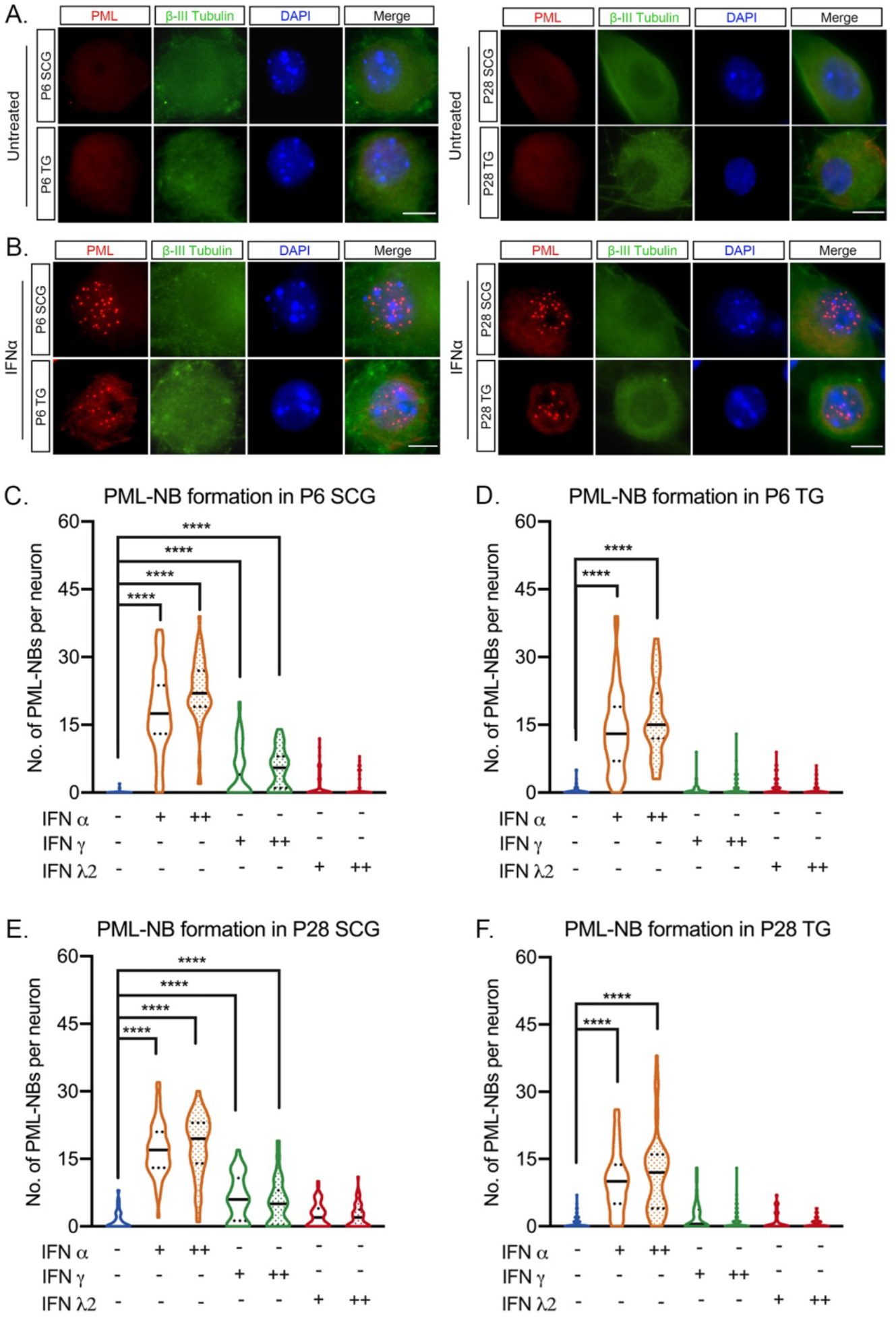
Type I IFN induces the formation of PML-NBs in primary peripheral neurons. (**A**) Representative images of primary neurons isolated from superior cervical ganglia (SCG) and sensory trigeminal ganglia (TG) of postnatal (P6) and adult (P28) mice stained for PML and the neuronal marker BIII-tubulin. (**B**) SCG and TG neurons isolated from P6 and P28 mice were treated with interferon (IFN)α (600 IU/ml) for 18h and stained for PML and BIII-tubulin. (**C-F**) Quantification of PML puncta in P6 and P28 neurons following 18h treatment with IFNα (150 IU/ml, 600 IU/ml), IFNγ (150 IU/ml, 500 IU/ml) and IFNλ2 (100 ng/ml, 500 ng/ml). Statistical comparisons were made using one way ANOVA with a Tukey’s multiple comparison (^****^ P<0.0001). Scale bar, 20μm.

In certain cell types, the transcription of certain PML-NB associated proteins, including PML, can be induced by either type I or type II interferon (IFN) treatment, which is correlated with an increase in PML-NB size and/or number per cell (Chelbi-Alix et al., 1995, Stadler et al., 1995). Therefore, we were interested in determining whether exposure of primary sensory or sympathetic neurons to different types of IFN resulted in PML-NB formation. Type I IFN treatment using IFN-alpha (IFNα) or IFN-beta (Fig. S1A) led to a significant induction of PML-NBs in both sensory and sympathetic neurons isolated from postnatal and adult mice. Representative images of IFNα-treated neurons are shown (Fig. 1B) and number of PML-NBs per neurons are quantified (Fig. 1C-1F). The increase in PML-NBs was comparable for both 150 IU/ml and 600 IU/ml of IFNα. Type II IFN (IFNγ) led to a more variable response with a small but significant increase in PML-NBs in a subpopulation of sympathetic neurons. However, IFNγ treatment of sensory neurons did not result in the formation of PML-NBs. Exposure of neurons to IFN-lambda 2 (IFN-λ2), a type III IFN, did not induce the formation of PML-NBs in either sympathetic or sensory neuron cultures (Fig. 1C-1F; Fig. S1B). Therefore, primary sympathetic and sensory neurons are largely devoid of PML-NBs but can form bodies upon exposure to type I IFNs.

The absence of PML-NBs in untreated primary neurons prompted us to investigate other known components of PML-NBs. We were particularly interested in ATRX and Daxx because like PML they have previously been found to be involved in restricting HSV lytic replication in non-neuronal cells (McFarlane et al., 2019, Alandijany et al., 2018, Lukashchuk and Everett, 2010, Cabral et al., 2018). Therefore, we investigated the localization of ATRX and Daxx in primary peripheral neurons. ATRX is a multifunctional, heterochromatin associated protein that is localized to PML-NBs in human and mouse mitotic cells and is largely characterized as interacting with the Daxx histone chaperone (Clynes et al., 2013, Lewis et al., 2010). In untreated neurons, we observed abundant ATRX staining throughout the nucleus in regions that also stained strongly with Hoechst (Fig. S1C, D). This potential co-localization of ATRX with regions of dense chromatin is consistent with a previous study demonstrating that in neurons ATRX binds certain regions of the cellular genome associated with the constitutive heterochromatin modification H3K9me3 (Noh et al., 2015). Importantly, this distribution of ATRX differs from what is seen in murine dermal fibroblasts (Fig. S1C, D) and other non-neuronal cells, where there is a high degree of colocalization between ATRX and PML (Alandijany et al., 2018). Following treatment with IFNα, we found a redistribution of ATRX staining and colocalization between ATRX and the formed PML-NBs, but the majority of ATRX staining remained outside the context of PML-NBs (Fig. S1C, D). Similar to PML, sympathetic SCG and sensory TG neurons isolated from both postnatal and adult mice were devoid of discreate puncta of Daxx staining (Fig. S1D), and we did not observe extensive Daxx staining in untreated neurons as we did for ATRX. We were unable to directly co-stain for Daxx and PML; however, treatment of neurons with IFNα did induce punctate Daxx staining that strongly colocalized with puncta of ATRX (Fig. S1D), which given our previous observation of ATRX co-localization with PML following type I IFN treatment we used as a correlate for PML-NBs. Therefore, PML-NBs containing their well characterized associated proteins are not present in cultured primary neurons but form in response to type I IFN exposure.

### Type I IFN treatment specifically at time of infection restricts reactivation of HSV-1 from primary sympathetic neurons without affecting initial infectivity or LAT expression

Because we observed that primary SCG neurons are largely devoid of PML-NBs and that PML-NBs form upon treatment with type I IFN treatment, we first wanted to clarify that latency was maintained in the absence of IFN and presumably without PML-NB formation, consistent with our previous data (Cuddy et al., 2020). SCG neurons were infected at a multiplicity of infection (MOI) of 7.5 plaque forming units (PFU)/cell with HSV-1 Us11-GFP presence of acyclovir (ACV). The ACV was removed after 6 days and the neuronal cultures were monitored to ensure the no GFP-positive neurons were present (Fig. 2A). We found that latency could be established and maintained for up to 5 days following removal of ACV (Fig. 2B). Reactivation was triggered by PI3K inhibition using LY294002, as previously described (Cliffe et al., 2015, Camarena, 2011, Kim et al., 2012, Kobayashi et al., 2012), and quantified based on the number of Us11- GFP neurons in the presence of WAY-150138 which blocks packaging of progeny genomes and thus cell-to-cell spread (Cliffe et al., 2015, van Zeijl et al., 2000). These data therefore indicate that exogenous IFN is not required to induce a latent state in this model system.

**Fig. 2.**
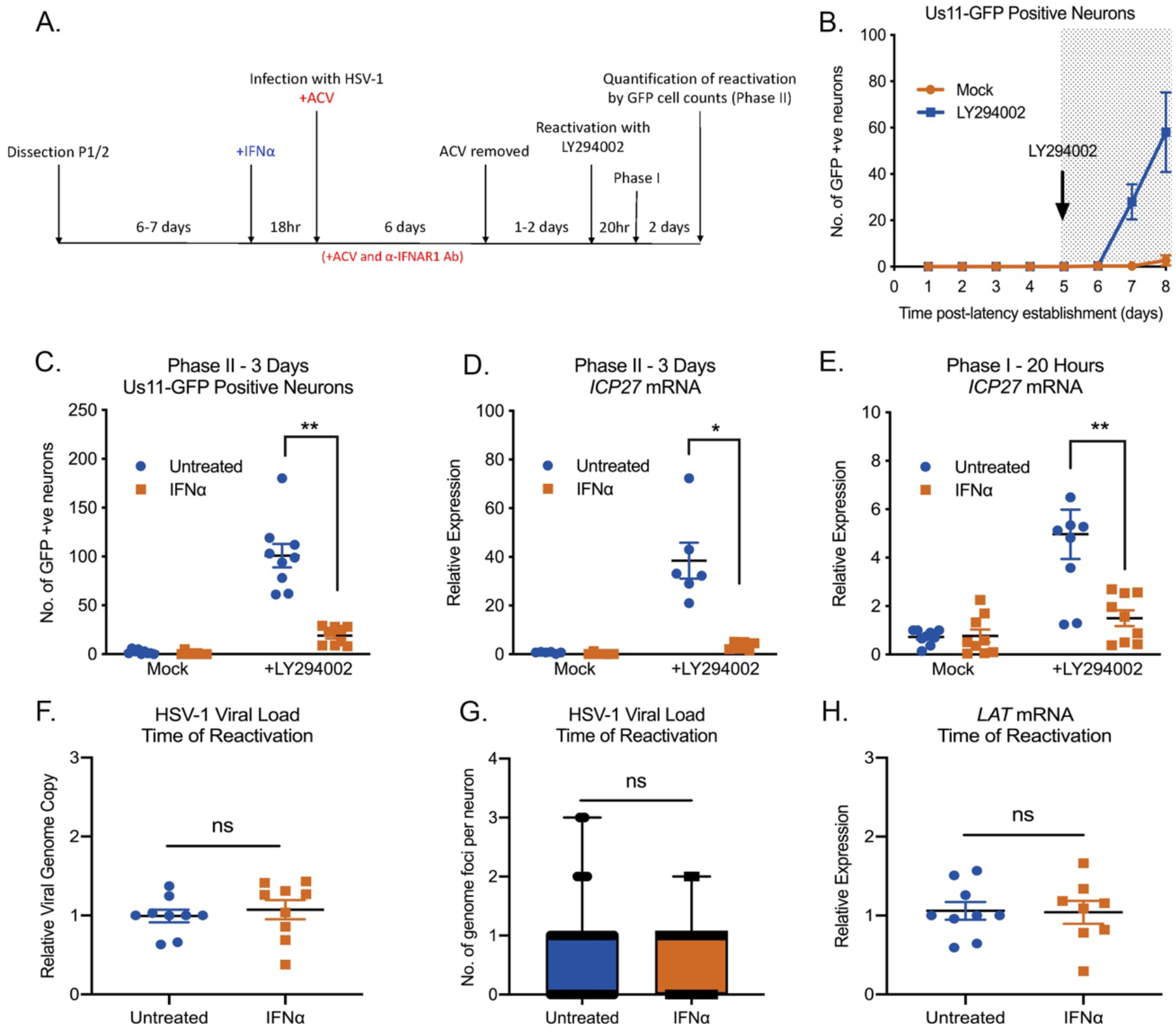
Type I IFN treatment solely at time of infection inhibits LY294002-mediated reactivation of HSV-1 in primary sympathetic SCG neurons. (**A**) Schematic of the primary postnatal sympathetic neuron-derived model of HSV-1 latency. (**B**) Reactivation from latency is quantified by Us11-GFP expressing neurons following addition of the PI3K inhibitor LY294002 (20μM) in the presence of WAY-150168, which prevents cell-to-cell spread. (**C**) Number of Us11-GFP expressing P6 SCG neurons infected with HSV-1 in the presence or absence of IFNα (600 IU/ml), then treated with an α-IFNAR1 neutralizing antibody. (**D**) RT-qPCR for viral mRNA transcripts at 3 days post- reactivation of SCGs infected with HSV-1 in the presence or absence of IFNα. (**E**) RT-qPCR for viral mRNA transcripts at 20 hours post-reactivation in SCGs infected with HSV-1 in the presence of absence of IFNα. (**F**) Relative amount of viral DNA at time of reactivation (8dpi) in SCG neurons infected with HSV-1 in the presence or absence of IFNα (600 IU/ml). (**G**) Quantification of vDNA foci detected by click chemistry at time of reactivation (8dpi) in SCG neurons infected with HSV-1 in the presence or absence of IFNα (600 IU/ml). (**H**) *LAT* mRNA expression at time of reactivation (8dpi) in neurons infected with HSV-1 in the presence or absence of IFNα (600 IU/ml). Statistical comparisons were made using Wilcoxon signed-rank test (ns not significant, ^*^ P<0.05, ^**^ P<0.01).

We next turned our attention to whether type I IFN treatment at the time of infection impacted the ability of HSV to establish latency or reactivate in this model system. SCG neurons were pre-treated with IFNα (600 IU/ml) for 18h and during the initial 2h HSV inoculation. Following inoculation, IFNα was washed out and an IFNAR1 blocking antibody was used to prevent subsequent type I IFN signaling through the receptor. Reactivation was induced and initially quantified based on the number of GFP positive neurons at 3-days post-stimuli. We found that full reactivation was restricted in neurons exposed to type I IFN just prior to and during *de novo* infection (Fig. 2C). We further confirmed this IFNα-mediated restriction of latency by the induction of lytic mRNAs upon reactivation. IFNα treatment at the time of infection significantly decreased the expression of immediate early gene (ICP27), early gene (ICP8) and late gene (gC) at 3 days post-reactivation (Fig. 2D, S2A, B). There were very few GFP- positive neurons and little to no viral gene expression in mock reactivated controls, further indicating that latency can be established in the presence and absence of IFN.

Reactivation of HSV in this system proceeds over two phases. GFP-positive neurons is a readout for full reactivation or Phase II. However, we and others have observed an initial wave of lytic gene expression that occurs prior to and independently of viral DNA replication at around 20 hours post-stimulus, termed Phase I (Cliffe and Wilson, 2017, Kim et al., 2012, Du et al., 2011, Cliffe et al., 2015). Therefore, to determine if IFNα treatment at the time of infection restricted the Phase I wave of lytic we carried out RT-qPCR to detect representative immediate-early (ICP27), early (ICP8), and late (gC) transcripts at 20 hours post addition of LY294002. We found significantly decreased expression in the IFNα-treated neurons (Fig. 2E, S2C, D). Therefore, type I IFN treatment solely at the time of infection has a long-term effect on the ability of HSV to initiate lytic gene expression and undergo reactivation.

Because IFN treatment could reduce nuclear trafficking of viral capsids during initial infection or impact infection efficiency, we next determined whether equivalent numbers of viral genomes were present in the neuronal cultures. At 8dpi, we measured relative viral DNA genome copy numbers in SCG neurons that were treated with IFNα compared to untreated controls and found no significant difference (Fig 2F). To further confirm that equivalent genomes were present in the neuronal nuclei, we infected neurons with HSV-1 containing EdC-incorporated genomes and performed click chemistry to detect vDNA foci. At 8 dpi, we found no significant difference in the average number of vDNA foci per nucleus of neurons treated with IFNα at the time of initial infection compared to untreated controls (Fig. 2G). Therefore, the restricted reactivation phenotype mediated by IFNα was not due to a decrease in the number of latent viral genomes.

The decreased reactivation observed with IFNα treatment could be secondary to changes in expression of the LAT and/or directly as a result of decreased viral genome accessibility. The HSV LAT, one of the only highly expressed gene products during latent infection, has been shown to modulate several features of latency, including the viral chromatin structure, lytic gene expression, and neuronal survival, as well as the efficiency of latency establishment and reactivation (Knipe and Cliffe, 2008, Cliffe et al., 2009, Chen et al., 1997, Thompson and Sawtell, 2001, Thompson and Sawtell, 1997, Gordon et al., 1995, Branco and Fraser, 2005). Therefore, the ability of HSV to undergo reactivation could be due to changes in LAT expression following IFNα treatment. However, when we evaluated LAT expression levels at 8 dpi by RT-qPCR, we found no difference between IFNα-treated and untreated neurons. This suggests that the IFNα- mediated restriction in reactivation does not appear to occur as a result of changes in expression of the LAT (Fig. 2H). Therefore, it is possible that the type I IFN-mediated restriction of HSV latency is due to changes to the latent genome that results in a decreased ability to undergo reactivation following PI3-kinase inhibition.

### Primary neurons have a memory of prior IFNα exposure characterized by persistence of PML-NBs

Because we observed a restriction in the ability of HSV to reactivate that occurred 7-8 days following type I IFN exposure, we went on to examine any long-term changes resulting from IFNα exposure. First, we investigated the kinetics of representative ISG expression. As expected, we saw a robust induction of *Isg15* and *Irf7* in IFNα-treated (600 IU/ml) neurons that persisted for at least 42 hours post-treatment post-addition of IFNα (this represents 1-day post-infection (dpi)). However, by 8 dpi, the time at which neurons were induced to reactivate, there was no difference in *Isg15* or *Irf7* expression in IFNα treated neurons vs untreated controls (Fig. 3A, B), indicating that these representative ISGs were not elevated at the time of reactivation. We also found no difference in *Isg15* or *Irf7* expression in HSV-1 infected neurons compared to uninfected controls in either the presence or absence of IFNα, suggesting that HSV-1 infection was not impacting IFN signaling pathways at a population level. PML has been previously characterized as an ISG product in non-neuronal cells (Chelbi-Alix et al., 1995, Stadler et al., 1995), and we found an approximate 5-fold-increased expression of *Pml* in primary sympathetic neurons following IFNα treatment which was less than the increased expression of *Irf7* and *Isg15* (approximate 350-fold- and 100-fold-increased expression respectively). *Pml* expression returned to untreated levels by 1 dpi (Fig. 3C).

**Fig. 3.**
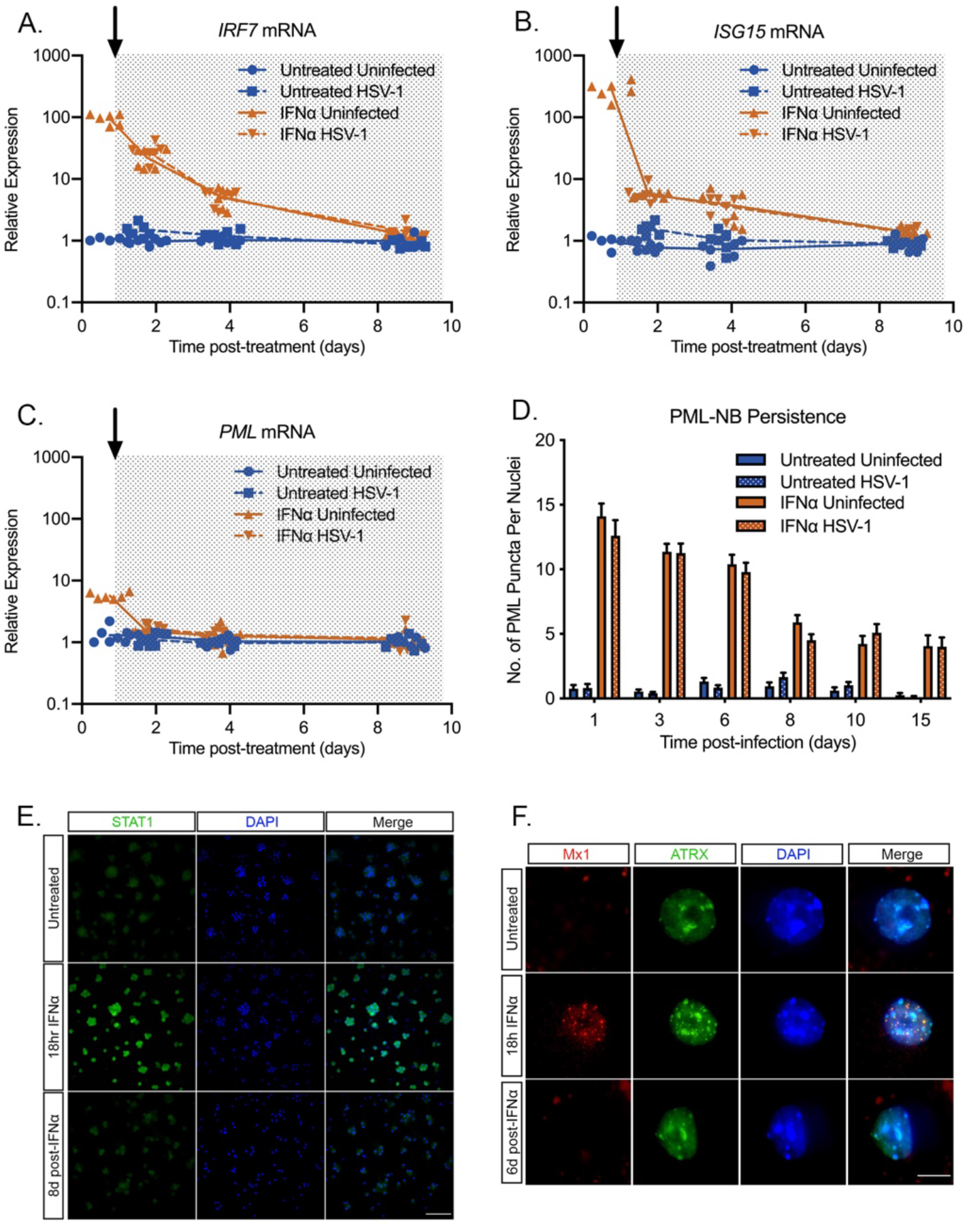
Type I IFN-induced PML-NBs persist in primary sympathetic neurons despite resolution of IFN signaling. (**A-C**) Kinetics of *ISG15, IRF7* and *PML* mRNA expression at 0.75, 1.75, 3.75 and 8.75 days post-IFNα (600 IU/ml) treatment. Arrow indicates the time of HSV-1 infection at 18hr post-interferon treatment. (**D**) Quantification of PML puncta at 1-, 3-, 6-, 8-, 10- and 15-days post-infection with HSV-1 in untreated and IFNα (600 IU/ml)-treated SCG neurons. (**E**) Representative images of P6 SCG neurons treated with IFNα (600 IU/ml) and stained for STAT1 at 18 hours and 8 days post-treatment. Scale bar, 100μm. (F) Representative images of P6 SCG neurons treated with IFNα (600 IU/ml) and stained for Mx1 at 18 hours and 6 days post-treatment. Scale bar, 20μm.

Although we did not detect maintained induction of IFN stimulated gene expression including *Pml*, we were intrigued as to whether PML-NBs persisted throughout the course of infection. To assess this, we first established whether PML-NBs persist even in the absence of sustained ISG expression. Quantifying the number of PML-NBs following IFNα (600 IU/ml) treatment, we found that the number of bodies remain elevated through 15 days post-treatment (Fig. 3D). We went on to investigate additional products of ISGs including STAT1 and Mx1 because of the availability of specific antibodies against these proteins. We observed robust STAT1 staining following IFNα exposure for 18 hours. However, by 8 days post infection we could not detect STAT1 staining in primary neurons indicating that synthesis of this IFNα-induced protein had returned to baseline (Fig. 3E). Similarly, we found induction of punctate Mx1 staining in neurons exposed to IFNα for 18 hours that was undetectable by day 6 post-treatment (Fig. 3F). Therefore, exposure of primary neurons to type I IFN led to a modest induction of *Pml* mRNA but resulted in long-term persistence of PML-NBs, even in the absence of continued IFN signaling and when antiviral protein products of other ISGs were undetectable.

### PML-NBs Persist and Stably Entrap Latent HSV-1 Genomes only if IFNα is Present at the Time of Initial Infection

The persistence of PML-NBs following IFN exposure raised the possibility that viral genomes are maintained within PML-NBs only in type I IFN-treated neurons. This would also suggest that PML-NB-associated genomes are less permissive for reactivation and provide us with an experimental system to investigate the contribution of PML-NBs to the maintenance of HSV latency. To determine whether viral genomes localize with PML-NBs in type I IFN-treated neurons, SCG neurons were pretreated with IFNα (600 IU/ml) then infected with HSV-1^EdC^ at an MOI of 5 PFU/cell in the presence of ACV and IFNα as described above. By co-staining for PML, we found that a large proportion of vDNA foci colocalized with PML-NBs in the IFNα-treated neurons over the course of infection. In untreated neurons that are largely devoid PML-NBs, very few genomes were colocalized to PML puncta as expected. Representative images are shown (Fig. 4A) and the percent of genome foci colocalized to PML-NBs is quantified (Fig. 4B). Furthermore, high-resolution Airy scan-based 3D confocal microscopy of IFNα-treated neurons revealed that vDNA foci were entrapped within PML-NBs (Fig. 4C, D), as has also been reported upon lytic infection of non-neuronal cell lines (Alandijany et al., 2018) and in latently infected TG *in vivo* (Catez et al., 2012). Rapid colocalization of viral DNA by PML-NBs during lytic HSV-1 infection of human fibroblasts occurs independently of type I IFN exposure, and we confirmed this was also true in dermal fibroblasts isolated from postnatal mice (Fig. S3A). Therefore, the presence of IFNα during initial infection can impact the long-term subnuclear localization of latent viral genomes in neurons by inducing PML-NBs that persist and stably entrap latent viral genomes.

**Fig. 4.**
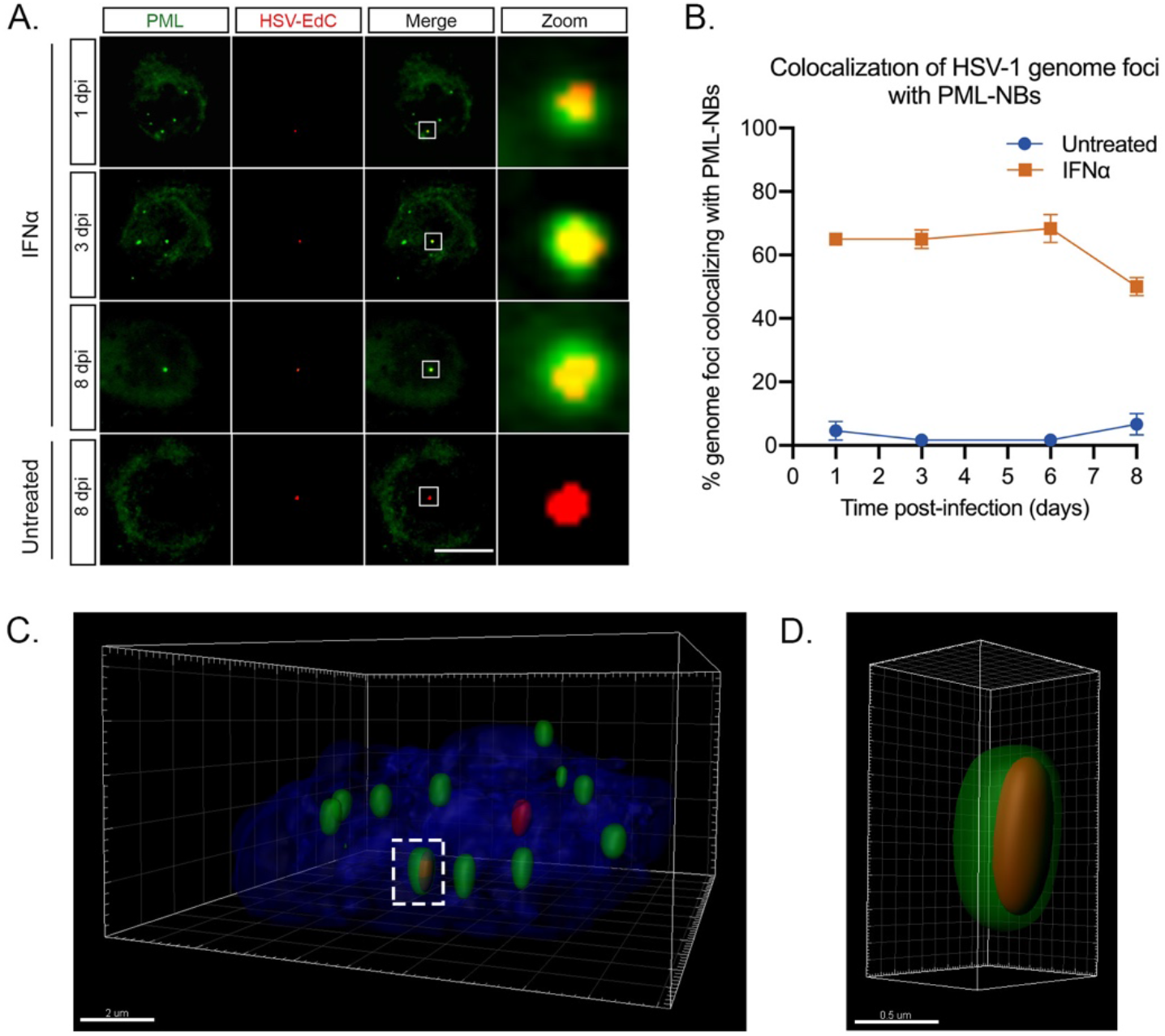
Type I IFN induced PML-NBs stably entrap vDNA throughout a latent HSV-1 infection of primary sympathetic neurons. (**A**) Representative images of vDNA foci detected by click chemistry to PML at 1, 3, 6 and 8 dpi in P6 SCG neurons infected with HSV-1^EdC^ in the presence or absence of IFNα (600 IU/ml). (**B**) Percent colocalization of vDNA foci detected by click chemistry to PML at 1, 3, 6 and 8 dpi in SCG neurons infected with HSV-1^EdC^ in the presence or absence of IFNα (600 IU/ml). Scale bar, 20μm. (**C**) 3D reconstruction of a high-resolution Z-series confocal image showing PML entrapment of a HSV-1^EdC^ vDNA foci. Scale bar, 2μm. (**D**) Enlargement of PML entrapped vDNA outlined by white dashed box. Scale bar, 0.5μm.

Thus far, our data indicate that the presence of IFNα during initial infection determines subnuclear positioning of latent viral genomes and the ability of genomes to reactivate in response to loss inhibition of PI3 kinase activity. We considered that type I IFN treatment could have a long-term effect on cell signaling pathways which could impact the ability of HSV to reactivate, so to determine the direct versus indirect effects on the viral genome itself, we next investigated whether the timing of IFNα exposure had a differential effect on the ability of viral genomes to reactivate. Therefore, we treated postnatal SCG neurons with IFNα (600 IU/ml) for 18h and during the 2h HSV inoculation (-18hpi) or exposed neurons to IFNα for 18h at 3 days prior to infection (- 3dpi). Following pretreatment at -3dpi, IFNα was washed out and an IFNAR1 blocking antibody was used. As expected, IFNα during initial infection significantly inhibited HSV reactivation, but surprisingly, IFNα treatment at -3dpi did not restrict reactivation as shown by the similar number of GFP-positive neurons at 72 hours post-reactivation when compared to untreated neurons (Fig. 5A). Surprisingly, we found that vDNA foci did not localize to PML-NBs in SCG neurons treated with IFNα at -3dpi (Fig. 5B). We confirmed that PML-NBs were present at the time of infection in neurons treated 3 day prior to infection (Fig. 5C), although we did detect slightly fewer PML-NBs per nucleus in neurons treated -3dpi compared to -18hpi (a mean of 17.57 versus 12.47 per nucleus respectively). We also confirmed comparable recruitment of known PML-NB-associated proteins ATRX and Daxx at 3 days post-IFNα treatment (Fig S4A, B). When IFNα treatment of SCG neurons is continued from -3pi through infection, or if SCG neurons treated at -3pi receive a second treatment of IFNα during infection, then a similar proportion of latent viral genomes colocalize with PML-NBs as with a single treatment during infection (Fig. S4C). This indicates type I IFN must be present during infection for vDNA to colocalize with IFN-induced PML-NBs.

**Fig. 5.**
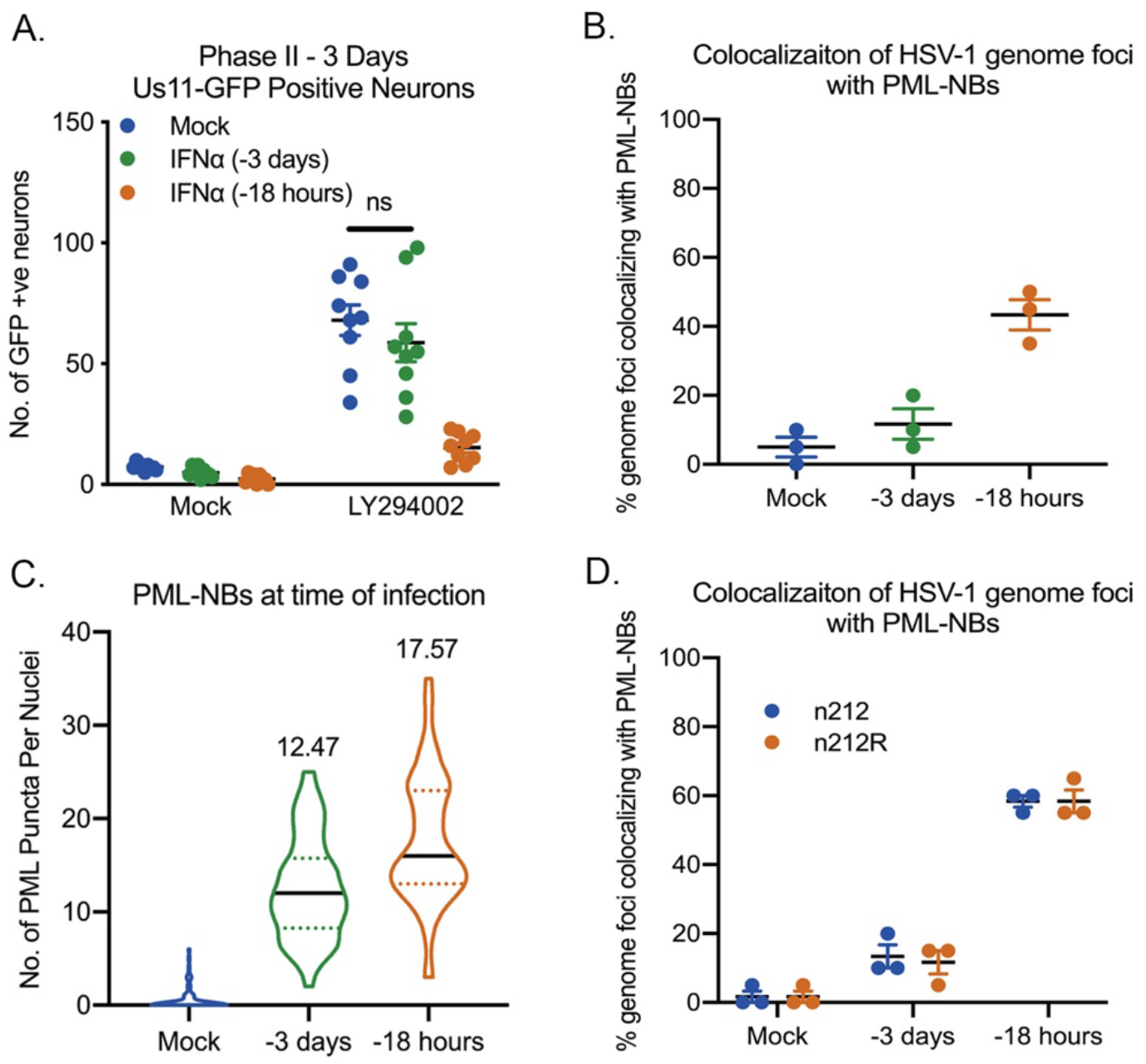
HSV-1 genomes only associate with PML-NBs when type I IFN is present during initial infection. (**A**) Number of Us11-GFP expressing P6 SCG neurons infected with HSV-1 following IFNα treatment for 18 hours prior to infection or for 18 hours at 3 days prior to infection. (**B**) Percent colocalization of vDNA foci detected by click chemistry to PML at 8 dpi in SCG neurons infected with HSV-1^EdC^ following IFNα treatment for 18 hours prior to infection or for 18 hours at 3 days prior to infection. (**C**) Quantification of PML puncta at time of infection in P6 SCG neurons treated with IFNα (600 IU/ml) for 18 hours prior to infection or for 18hours at 3 days prior to infection. (**D**) Percent colocalization of vDNA foci detected by click chemistry to PML at 3 dpi in SCG neurons with HSV-1^EdC^ infected with ICP0-null mutant HSV-1, n212, or a rescued HSV-1 virus, n212R, in P6 SCG neurons treated with IFNα for 18 hours prior to infection or for 18hours at 3 days prior to infection.

The HSV Infected Cell Protein 0 (ICP0) is a RING-finger E3 ubiquitin ligase that disrupts PML-NBs (Boutell et al., 2011, Boutell et al., 2002, Alandijany et al., 2018, Cuchet-Lourenco et al., 2012, Chelbi-Alix and de The, 1999, Muller et al., 1998, Everett et al., 1998) and known to be expressed during the establishment of latency (Cliffe et al., 2013). Therefore, the colocalization of latent viral genomes to PML-NBs and ultimately the ability of HSV to undergo reactivation could be due the presence of IFNα during initial infection and its effect on the localization or amount of ICP0. To investigate the distribution of ICP0 at early time points post-infection, SCG neurons were treated with IFNα at either -3dpi or -18hpi and infected at a MOI of 7.5 PFU/cell with HSV-1 Us11-GFP in the presence of acyclovir (ACV). In both treatment groups, ICP0 staining similarly colocalized with puncta of ATRX, a correlate for PML-NBs, at 3, 6 and 9 hours post-infection (Fig. S4D, E). To further investigate the effect of ICP0 on the colocalization of latent viral genomes to PML-NBs, we generated an EdC-labeled ICP0- null mutant strain (n212) and found that ICP0 had no impact on the ability of vDNA foci to colocalize to PML-NBs (Fig. 5D). Taken together, these data demonstrate that association of latent viral genomes with PML-NBs in peripheral neurons is dependent on the formation of type I IFN-induced PML-NBs and the presence of type I IFN during initial infection and is independent of ICP0 expression.

### PML is Required for the IFNα-dependent Restriction of HSV-1 Latency

To determine whether the stable association of viral genomes with PML-NBs directly contributes to the IFNα-dependent restriction of HSV reactivation, we investigated whether PML depletion was sufficient to restore the ability of the latent viral genomes to reactivate. A previous study has shown that PML-dependent recruitment of HIRA to ISG promoters contributes to the up-regulation of gene expression as a result of cytokine release in response to HSV infection (McFarlane et al., 2019). Although carried out in non-neuronal cells, this study and others (Ulbricht et al., 2012, Kim and Ahn, 2015, Scherer et al., 2016, Chen et al., 2015) suggest that PML itself may contribute to ISG upregulation, so to determine whether PML was indeed required for ISG stimulation in SCG neurons we carried out RNA sequence analysis in IFNα-treated neurons depleted of PML. Postnatal SCG neurons were transduced with lentiviral vectors expressing non-targeting control or PML-targeting shRNAs (shCtrl and shPML, respectively) and then mock treated or treated with IFNα (600 IU/ml) for 18h prior to RNA extraction for next generation sequencing. High confidence reads were used for gene expression and gene ontology (GO) analysis. As expected, treatment of shCtrl transduced neurons with IFNα caused large changes in differentially regulated gene expression, with an enrichment of upregulated genes involved in immune system regulation. Similar to control neurons, PML depleted neurons also significantly upregulated the expression of genes involved in the response to IFNα stimulation. We found that of the total of 248 genes upregulated >1.5-fold following IFNα treatment, 83.47% of these genes were shared between the shCtrl- and shPML-treated groups (Fig. 6A). Furthermore, we found similar ISG expression (Fig. 6B) and GO pathway enrichment (Fig. 6C). Therefore, in primary SCG neurons, the expression of ISGs in response to exogenous IFNα is largely independent of PML expression.

**Fig. 6.**
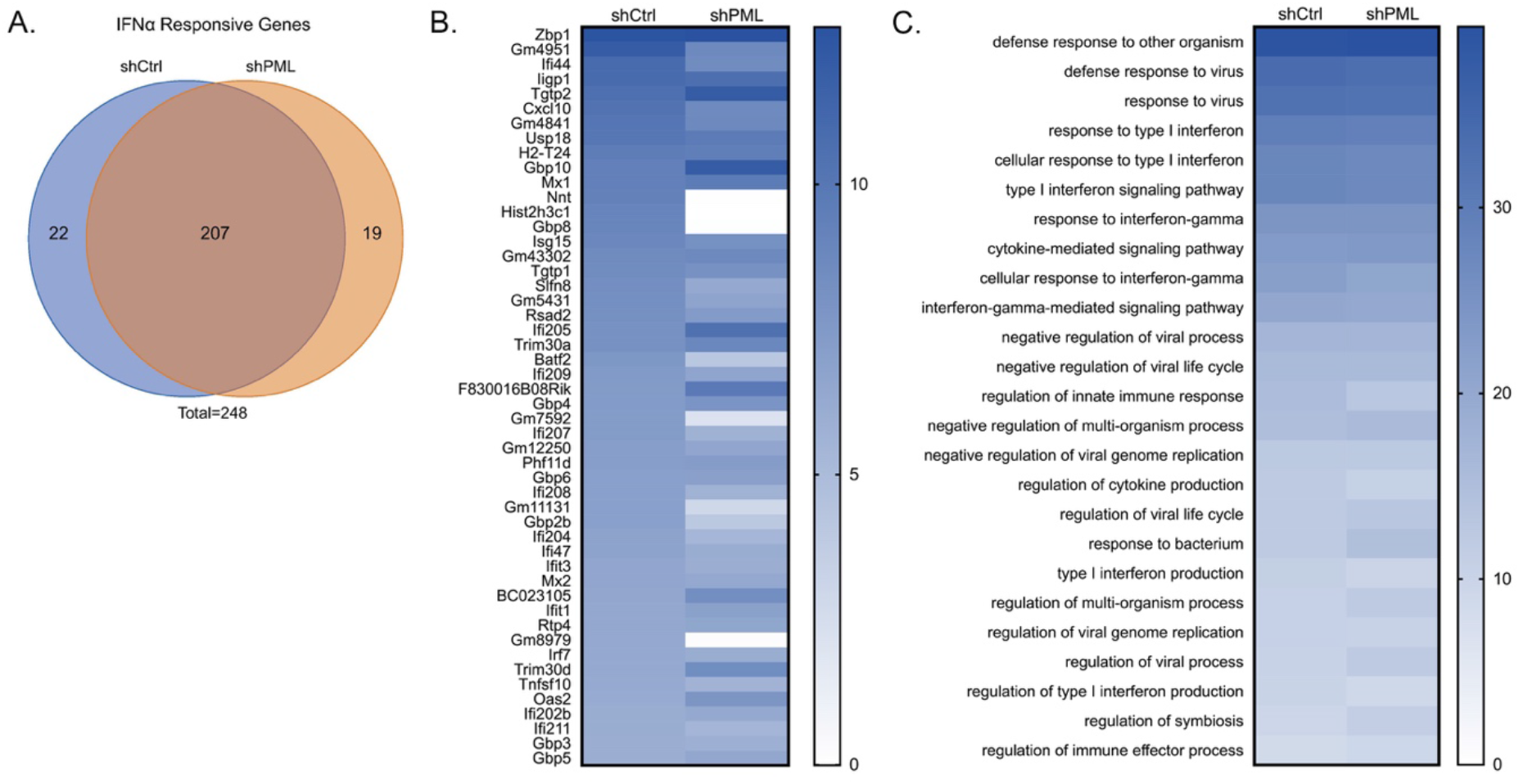
PML is not required for ISG induction in primary postnatal sympathetic neurons. (**A**) P6 SCG neurons were transduced with either control non-targeting shRNA or shRNA targeting PML for 3 days, then treated with IFNα (600. IU/ml) for 18 hours. Total genes >1.5-fold higher in IFNα (600 IU/ml) treated cells than untreated cells were subdivided into three groups: shCtrl-treated neurons only. shPML-treated neurons only. Both shCtrl and shPML neurons (Shared). (**B**) Gene expression heat map of top 50 most upregulated genes in P6 neurons transduced with control non- targeting shRNA or shRNA targeting PML for 3 days, then treated with IFNα (600 IU/ml) for 18 hours. (**C**) Heat map of the top 25 shared upregulated GO terms in P6 neurons transduced with control non-targeting shRNA or shRNA targeting PML for 3 days, then treated with IFNα for 18 hours.

Because PML depletion did not prevent the induction of type I IFN response genes in SCG neurons, we were able to examine the effect of PML depletion prior to infection on the IFNα-mediated restriction of HSV-1 reactivation. SCG neurons were transduced with lentiviral vectors expressing different PML-targeting shRNA or control non-targeting shRNA. PML depletion was confirmed by average number of PML-NBs per nucleus in neurons transduced for 3 days then treated with IFNα (Fig. 7A). As expected, we found a significant decrease in the percent of vDNA foci stably colocalizing with PML-NBs at 8 dpi in the shPML-treated neurons compared to shCtrl-treated neurons (Fig. 7B). Furthermore, we assessed reactivation in neurons infected with HSV-1 in the presence or absence of IFNα (150 IU/ml) at 3 days post-transduction. In these experiments, neurons were infected with a Us11-GFP gH-null virus, which is defective in cell-to-cell spread and eliminates the need for WAY-150138 during reactivation. In untreated neurons, we found no difference in reactivation (Fig. 7C, D). In addition, PML depletion had no effect on the number of GFP-positive neurons in the non-reactivated samples, indicating that in this system that PML was not required for the establishment of latency. However, in neurons treated with IFNα at the time of initial infection, depletion of PML using either of the three PML shRNAs significantly increased the ability of HSV to reactivate (Fig. 7E, F). Moreover, there was no significant difference between the PML depleted, IFNα-treated neurons and the non-IFNα treated neurons, indicating that PML depletion fully restored the ability of HSV to reactivate from type I IFN treated neurons. Taken together, these data demonstrate that type I IFN exposure solely at the time of infection results in entrapment of viral genomes in PML-NBs to directly promote a deeper form of latency that is restricted for reactivation.

**Fig. 7.**
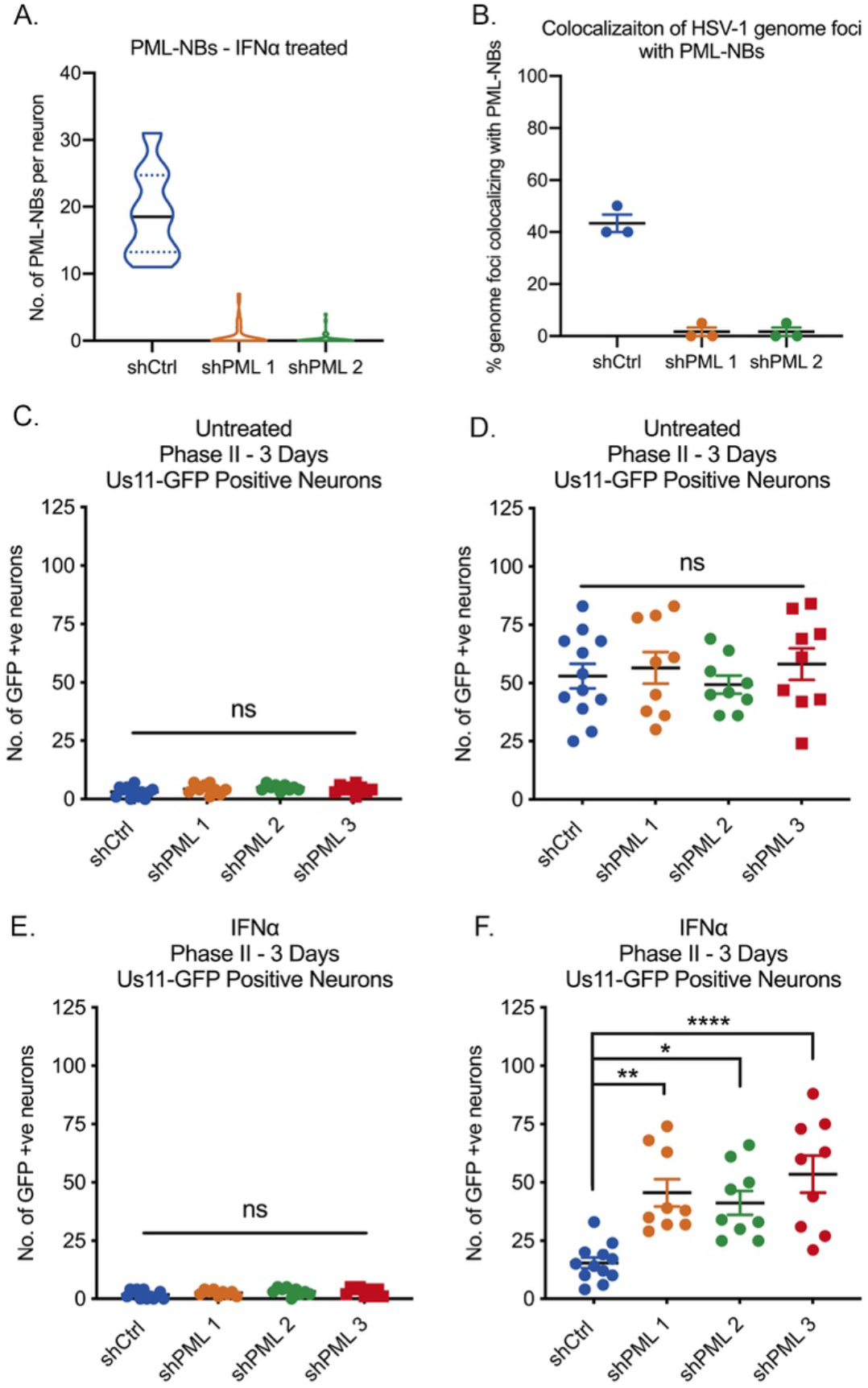
Depletion of PML with shRNA-mediated knockdown prior to infection restores HSV-1 reactivation in type I interferon-treated primary sympathetic neurons. (**A**) Quantification of PML puncta in P6 SCG neurons transduced with either control non-targeting shRNA or shRNA targeting PML for 3 days, then treated with IFNα (600 IU/ml) for 18 hours. (**B**) Quantification of colocalization of vDNA foci detected by click chemistry to PML in primary SCG neurons transduced with shRNA targeting PML for 3 days prior to being infected with HSV-1^EdC^ in the presence or absence of IFNα (150 IU/ml). (**C-F**) Number of Us11-GFP expressing P6 SCG transduced with shRNA targeting PML for 3 days prior to infection with HSV-1 in the presence or absence of IFNα (150 IU/ml). Statistical comparisons were made using one way ANOVA with a Tukey’s multiple comparison (ns not significant, ^*^ P<0.05, ^**^ P<0.01, ^****^ P<.0001).

### Depletion of PML After the Establishment of Latency Enhances Reactivation in IFNα- treated Neurons

To explore the long-term effect of stable PML-NB-association on the latent viral genome, we next tested whether PML depletion after the establishment of latency was sufficient to restore the ability of the latent viral genomes to reactivate following treatment with a physiological stimulus of reactivation. In these experiments, neurons were infected with Us11-GFP gH null HSV-1 virus in the presence or absence of IFNα (150 IU/ml) and subsequently transduced with lentiviral vectors expressing PML-targeting shRNA or control non-targeting shRNA at 1 dpi. Under these experimental conditions, PML knockdown post-infection significantly increased the ability of HSV to reactivate from IFNα treated neurons but not untreated neurons, albeit reactivation was not restored to levels seen in untreated neurons (Fig. 8A-D). As expected, we found that only a small proportion of vDNA foci stably colocalize with PML-NBs at 8 dpi in the shPML-treated neurons compared to vDNA foci in the shCtrl-treated neurons. Representative images are shown (Fig. 8E) and the percent of genome foci colocalized to PML-NBs is quantified (Fig. 8F). Therefore, PML depletion post-infection does not result in spontaneous reactivation of PML-NB-associated viral genomes, indicating that they are still in a repressed state and/or lack the necessary factors required to initiate gene expression. However, depletion of PML does partially restore the ability of HSV to enter the lytic from IFN-treated neurons in response to a reactivation stimulus.

**Fig. 8.**
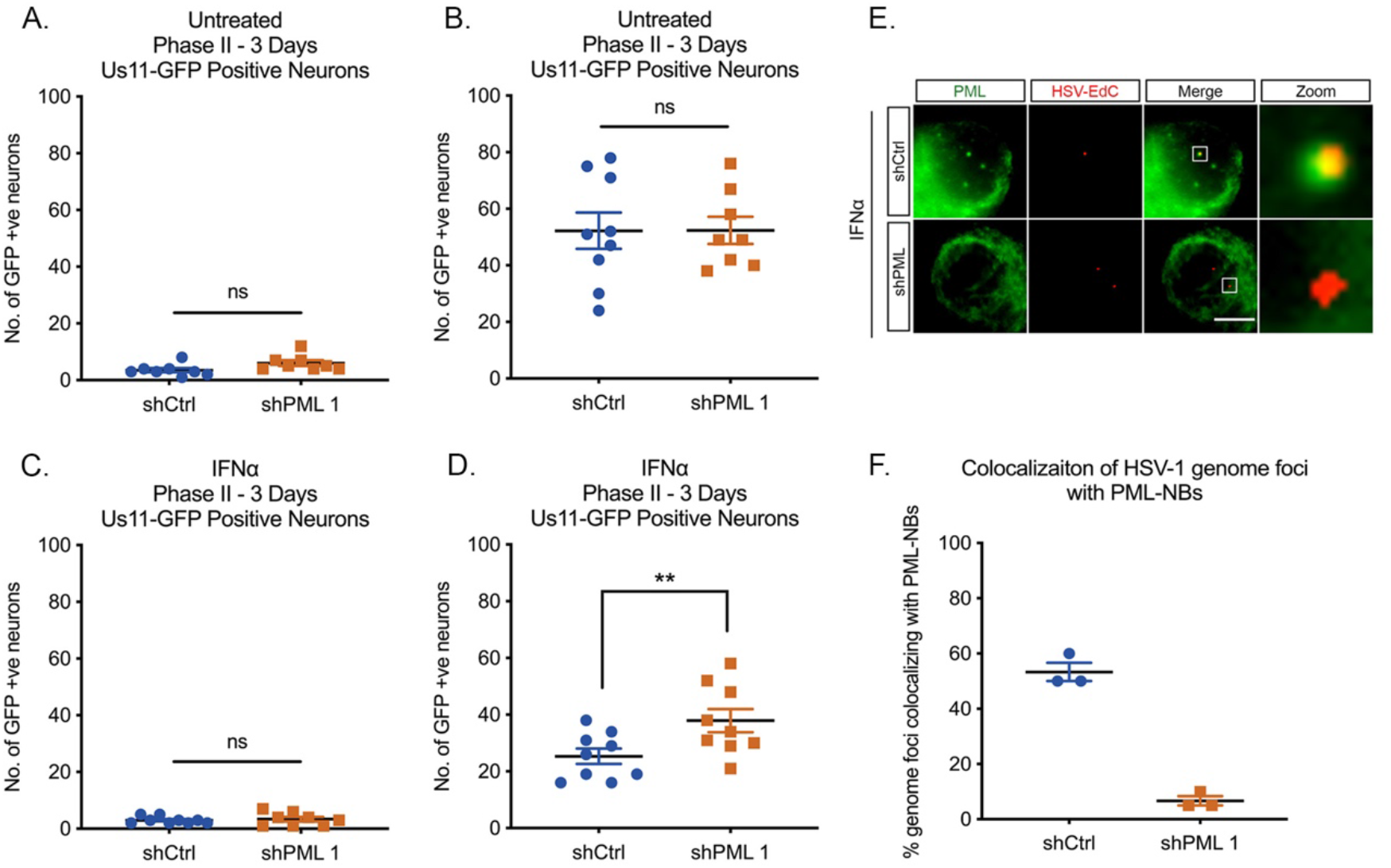
Depletion of PML with shRNA mediated knockdown post-infection partially restores HSV-1 reactivation in type I interferon-treated primary sympathetic neurons. (**A-D**) Number of Us11-GFP expressing P6 SCG transduced with shRNA targeting PML at 1 day post-infection with HSV-1 in the presence or absence of IFNα (150 IU/ml). (**E**) Representative images of vDNA foci detected by click chemistry 8 dpi in P6 SCG neurons transduced with shRNA targeting PML at 1 day post- infection with HSV-1 in the presence or absence of IFNα (150 IU/ml). (**F**) Quantification of colocalization of vDNA foci detected by click chemistry to PML in primary SCG neurons transduced with shRNA targeting PML at 1 day post-infection with HSV-1^EdC^ in the presence or absence of IFNα (150 IU/ml). Statistical comparisons were made using one way ANOVA with a Wilcoxon signed-rank test (ns not significant, ^**^ P<0.01). Scale bar, 20μm.

## Discussion

The considerable heterogeneity observed at the neuronal level in the colocalization of viral genomes with different nuclear domains may reflect in different types of latency that are more or less susceptible to reactivation. The determinants of this heterogeneity and a direct link between the subnuclear localization of a latent genome and its ability to reactivate following a given stimulus was not known. Using a primary neuronal model of HSV latency and reactivation, we found that the presence of type I IFN solely at that time of initial infection acts as a key mediator of the subnuclear distribution of latent viral genomes in neurons and promotes a more restricted form of latency that is less capable of reactivation following disruption of NGF-signaling. Importantly, we show that activation of the type I IFN signaling pathway in peripheral neurons induces the formation of PML-NBs, which stably entrap a proportion of latent genomes. Importantly, we show that this IFN-dependent restriction is mediated by PML, suggesting that PML-NBs are directly responsible for the observed restriction of reactivation.

PML-NBs typically number 1-30 bodies per nucleus in non-neuronal cells (Bernardi and Pandolfi, 2007). In the mouse nervous system, however, PML mRNA expression levels have previously been found to be low as measured by *in situ* hybridization (Gray et al., 2004). PML protein is enriched in neural progenitor cells, but the induction of differentiation results in the downregulation of PML both at a transcriptional and protein level, and PML mRNA expression is undetectable in post-mitotic neurons in many regions of the developing brain (Regad et al., 2009). Our findings in postnatal peripheral neurons further support these observations. Interestingly, PML has been shown to be re-expressed in both adult mouse and human brains, but often PML-NBs are associated with intranuclear inclusions in the context of pathological conditions, such as Guillain-Barre syndrome (Hall et al., 2016, Woulfe et al., 2004, Villagra et al., 2004). In our study, we could not detect PML-NBs in adult primary neurons isolated from the SCG or the TG. In contrast to our findings, PML-NBs have previously been shown to be present in adult TG neurons (Catez et al., 2012, Maroui et al., 2016). However, Catez et al. (2012) describes a subpopulation of adult TG neurons that did not display any PML signal in the nucleus. In addition, adult TG neurons isolated from humans at autopsy may reflect neurons that had previously been exposed to type I IFNs. The functional significance of peripheral neurons lacking PML-NBs is unclear, but could be linked to the capacity of neurons to undergo dynamic rearrangement of local and global nuclear architecture during maturation or neuronal excitation. An absence of PML-NBs in neurons could also contribute to their resistance to apoptosis, as PML has also been shown to play a role in cell death through the induction of both p53-dependent and -independent apoptotic pathways (Guo et al., 2000, Wang et al., 1998, Quignon et al., 1998). Whether PML-mediated regulation of these pathways occurs in the context of PML-NBs or by PML itself is unclear, but interestingly, the pro-apoptotic functions of Daxx, a PML-NB-associated protein, may require localization to PML-NBs in certain cell types (Croxton et al., 2006). Furthermore, our *in vitro* model using pure populations of intact neurons is devoid of the immune responses and complexities of intact animals, and we cannot rule out the possibility that axotomy or the processing of the neurons *ex vivo* could lead to PML-NB disruption or dispersal. However, notwithstanding these caveats, primary neurons provide an excellent model system to understand the impact of extrinsic immune factors and PML-NBs to the altering the nature of HSV latency.

Peripheral neurons are capable of responding to type I IFN signaling, given the robust induction in ISG expression and formation of PML-NBs following treatment with IFNα, and this is supported by a number of previous studies (Yordy et al., 2012, Katzenell and Leib, 2016, Song et al., 2016, Barragan-Iglesias et al., 2020). Importantly, however, peripheral neurons produce little to no type I interferons upon HSV infection (Yordy et al., 2012, Rosato and Leib, 2014), indicating that IFN production arises from other surrounding infected cells. Infected fibroblasts at the body surface, as well as professional immune cells, have been shown to produce high levels of IFNα/β after HSV infection (Hochrein et al., 2004, Rasmussen et al., 2007, Rasmussen et al., 2009, Li et al., 2006). In addition, there is evidence of elevated type I IFN in peripheral ganglia during HSV-1 infection (Carr et al., 1998), suggesting that glial or immune cells located adjacent to peripheral neuron cell bodies are capable of type I IFN production. It will be important to delineate if the inflammatory environment at the initial site of infection acts on neuronal axons to prime the neuron for a more repressed latent infection or if inflammatory cytokines in the ganglia are crucial for promoting a more repressive state. Although responsive to IFN, primary peripheral and cortical mouse neurons have previously been shown to have inefficient type I IFN-mediated anti-viral protection compared to non-neuronal mitotic cells (Yordy et al., 2012, Kreit et al., 2014). One study showed that DRG neurons are less responsive to type I IFN signaling and used an absence of cell death upon IFN treatment as one of their criteria (Yordy et al., 2012). It should be noted that different cell types display specific responses to type I IFN signaling and peripheral neurons have even been reported to be more protected from cell death stimuli following IFN treatment (Chang et al., 1990). Our model of HSV-1 latency and reactivation in primary sympathetic neurons highlights a type I IFN response that is PML-dependent and suggests a role for neuronal IFN signaling in promoting a more restricted latent HSV-1 infection.

Prior to this study, it was not clear whether viral genomes associated with PML- NBs were capable of undergoing reactivation. In response to inhibition of NGF- signaling, our data demonstrate that PML-NB associated genomes are more restricted for reactivation given that 1) IFN induces PML-NB formation and increased association with viral genomes with PML-NBs, 2) IFN pretreatment promotes restriction of viral reactivation and 3) the ability of viral genomes to reactivate from IFN-treated neurons increases with PML knock-down either prior to or following infection. Previous work by Cohen et al. (2018) showed that quiescent genomes associated with PML-NBs in fibroblasts can be transcriptionally reactivated by induced expression of ICP0. However, this previous study did not address the capability of viral genomes to reactivate in the absence of viral lytic protein (i.e. during reactivation from latency in neurons). In a further study using primary neurons, treatment of quiescently-infected neurons with the histone deacetylase inhibitor, trichostatin A (TSA), could lead to disruption of PML-NBs and induce active viral transcription in a subset of PML-NB-associated genomes (Maroui et al., 2016). However, the mechanisms of reactivation following TSA treatment are not known, and may be direct via altering the HSV chromatin structure or indirect via increasing the acetylation levels of histones or non-histone proteins, including PML. How increased acetylation relates to the physiological triggers that induce HSV reactivation is not clear. In contrast, loss of neurotrophic signaling can occur in response to known physiological stimuli that trigger HSV reactivation (Suzich and Cliffe, 2018). Although we cannot rule out the possibility that different stimuli have the potential for PML-NB associated genomes to undergo reactivation, this study clearly demonstrates that at least one well characterized trigger of reactivation cannot efficiently induce PML-NB associated genomes to undergo transcription.

Our results identify a persistence of PML-NBs, an IFN-mediated innate immune response, that allows for long-term restriction of latent viral genomes in the absence of continued ISG expression. Interestingly, type I IFN-induced PML-NBs persisted for up to 15 days post-treatment both in the presence and absence of viral infection. Given the absence of PML-NBs in our untreated peripheral neurons, this induction and persistence could represent neuron-specific innate immune memory. The persistence of PML-NBs in neurons may alter the subsequent response to IFN and/or viral infection, and it will be interesting to determine whether there is trained immunity in neurons such that subsequent responses differ from the first exposure. What is clear from our results however is the role of PML and IFN exposure in sustained repression of the latent HSV genome. Even in the absence of known chromatin changes that occur on the PML associated viral genome, this long-term effect on the ability of the HSV-1 genome to respond to an exogenous signal and restriction of reactivation is reminiscence of the classical definition of an epigenetic change (of course in the case of post-mitotic neurons in the absence of inheritance).

PML-NBs are known to play a role in the restriction of viral gene expression in non-neuronal cells, but the potential mechanism of PML-NB-mediated HSV gene silencing in neurons is unknown. During latency, the viral genome is enriched with histone post-translational modifications (PTMs) consistent with repressive heterochromatin, including H3K9me2/3 and H3K27me3, and it is possible that PML-NBs play a role in the association of viral genomes with core histones, repressive PTMs or heterochromatin-associated proteins (Cliffe et al., 2009, Kwiatkowski et al., 2009, Wang et al., 2005). In a model of quiescence utilizing human primary fibroblasts and a replication deficient virus, HSV genomes associated with PML-NBs were almost exclusively enriched with the H3.3K9me3 chromatin mark (Cohen et al., 2018). Therefore, it is tempting to speculate that PML-associated latent genomes are specifically enriched for H3K9me3 and not H3K27me3. However, in a previous study, we have found that H3S10 becomes phosphorylated during transcriptional activation following a reactivation stimulus (Cliffe et al., 2015) and viral genomes co-localize with regions of H3K9me3S10p in neurons that were not pre-treated with IFN (Cuddy et al., 2020). In addition, removal of H3K9 methylation is required for HSV reactivation (Liang et al., 2013, Liang et al., 2009). Together, these studies suggest that H3K9me3 is present on reactivation component genomes. However, it may be that different combinations of modifications exist on reactivation competent versus repressive genomes or that PML-NB associated genomes are less accessible for reactivation due to physical compaction of the genome and/or association with different histone reader proteins. Going forward, the primary neuronal system provides an excellent model to delineate the specific epigenetic contributions of PML-NBs to promoting a more repressive form of HSV latency.

Although PML-NBs promote a more restricted form of latency, we have shown that latency can be established in the absence of IFN treatment and PML-NBs. Even in IFN-treated neurons, only a proportion of the latent viral genomes co-localized with PML-NBs. This indicates that latent genomes associate with other subnuclear regions and proteins that may promote the assembly and/or maintenance of repressive heterochromatic histone modifications. This supports previous observations that HSV-1 viral genomes also co-localize with centromeric repeats and other, undefined nuclear domains in latently infected TG *in vivo* (Catez et al., 2012). For example, the viral genome is known to be enriched for H3K27me3 (Cliffe et al., 2009, Kwiatkowski et al., 2009, Wang et al., 2005), which can be bound by Polycomb group proteins. Interestingly, we have found that the multi-functional, chromatin remodeler protein ATRX has abundant nuclear staining in neurons and, in contrast to non-neuronal cells, is localized outside of PML-NBs. ATRX staining overlapped with Hoechst DNA staining in our primary neurons, suggesting its localization with AT-rich heterochromatin regions (Bucevicius et al., 2019). ATRX has previously been shown to interact with a variety of proteins, including methyltransferases and other heterochromatin-associated proteins, to promote transcriptional repression (Clynes et al., 2013, Lewis et al., 2010, Noh et al., 2015), as well as target chromatin through direct interactions with specific histone posttranslational modifications (PTMs), including H3K9me3-containing peptides (Noh et al., 2015). ATRX can also act as a histone chaperone, forming a complex with the death domain associated protein (Daxx) to catalyze the deposition of histone H3.3 (Lewis et al., 2010). Although we saw only faint staining of Daxx in our primary neurons, the ATRX/Daxx complex has been shown to promote the initial repression of the infecting viral genomes in non-neuronal cells (Lukashchuk and Everett, 2010). Ultimately, the repressive PTMs on latent viral genomes are likely bound by ATRX, Polycomb group proteins or other repressive cellular proteins independently of PML-NBs. Investigating the identity, mechanism of targeting and role of these proteins in the induction and maintenance of latency will ultimately facilitate the development of antiviral therapeutics that target the latent stage of infection to prevent reactivation.

## Acknowledgements

We thank Dr. Ian Mohr at New York University for the gift of the Us11-GFP virus and Gary Cohen at the University of Pennsylvania for SCgHZ. This work was supported by R21AI151340 (ARC), R01NS105630 (ARC), NIH/NEI F30EY030397 (JBS), NIH/NIAID T32AI007046 (JBS and SRC), T32GM007267 (JBS), NIH/NIGMS T32GM008136 (SD) and MRC (https://mrc.ukri.org) MC_UU_12014/5 (CB).

---

**Supplemental Fig. 1.**
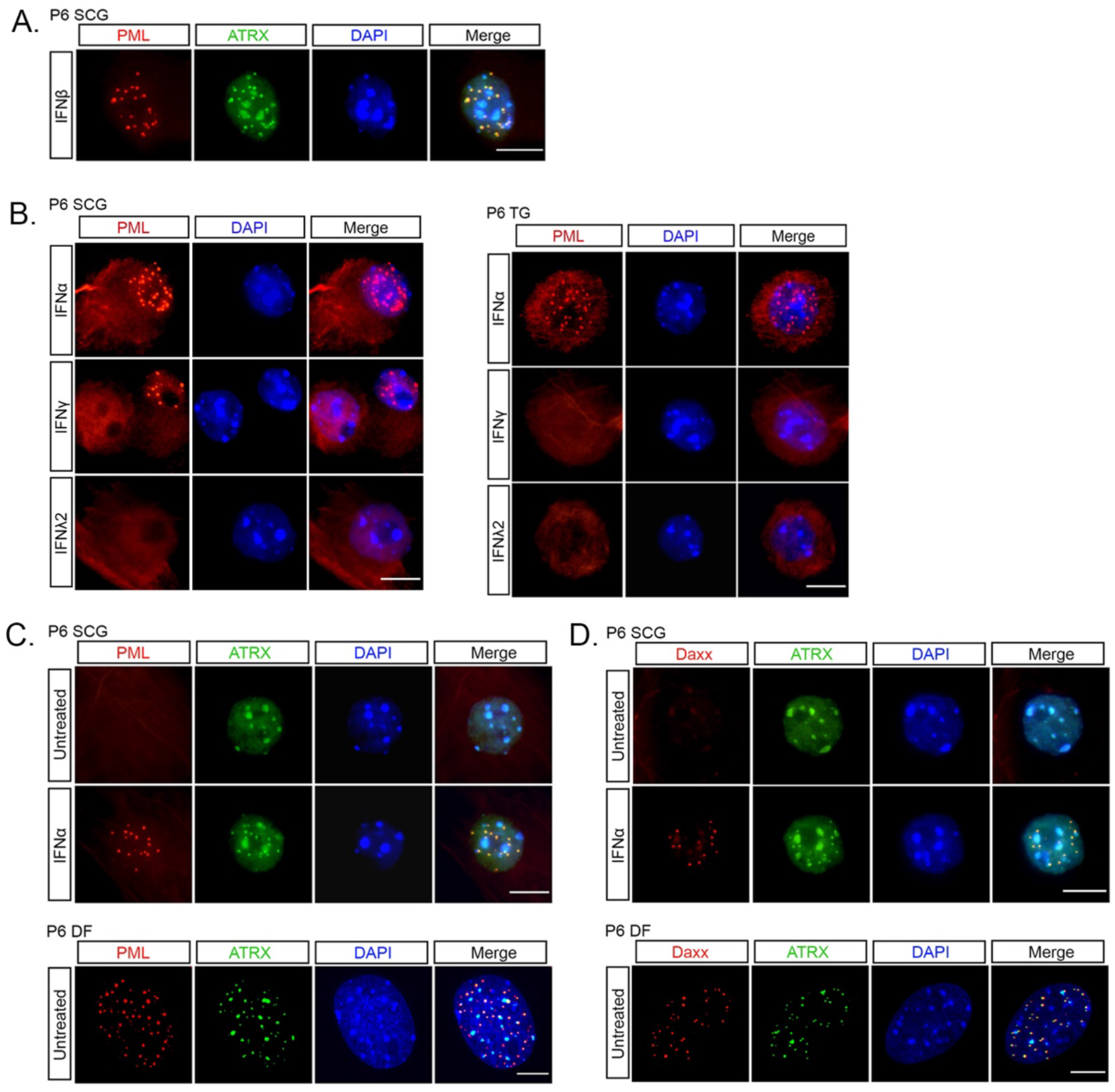
Type I IFN alters the sub-cellular localization of ATRX and Daxx in primary peripheral neurons. (**A**) Representative images of P6 SCG neurons treated with IFNβ (150 IU/ml) and stained for PML and ATRX. (**B**) Representative images of P6 SCG and TG treated with IFNα (600 IU/ml), IFNγ (500 IU/ml) and IFNλ2 (500ng/ml) and stained for PML. (**C**) Representative images of untreated or IFNα-treated P6 SCG neurons stained for PML and ATRX. (**D**) Representative images of untreated or IFNα-treated P6 SCG neurons stained for Daxx and ATRX. P6 dermal fibroblasts (DF) isolated from the same mice were used as a non-neuronal control. Scale bar, 20μm.

**Supplemental Fig. 2.**
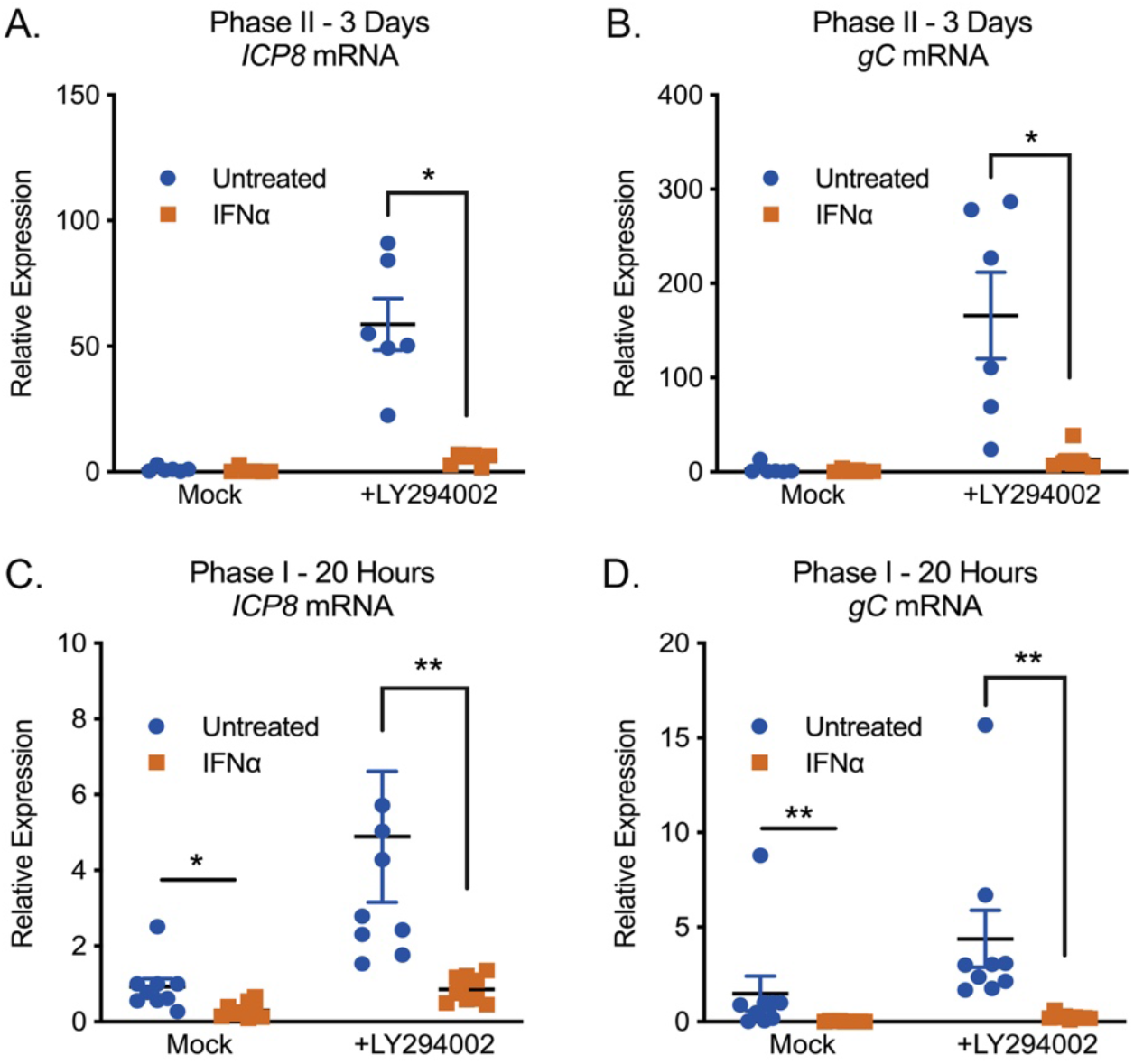
Type I IFN treatment solely at time of infection inhibits LY294002-mediated reactivation of HSV-1 in primary sympathetic SCG neurons. (**A**) RT-qPCR for viral mRNA transcripts at 3 days post-reactivation of SCGs infected with HSV-1 in the presence or absence of IFNα (600 IU/ml). (**B**) RT-qPCR for viral mRNA transcripts at 20 hours post-reactivation in SCGs infected with HSV-1 in the presence or absence of IFNα (600 IU/ml). Statistical comparisons were made using one way ANOVA with a Wilcoxon signed-rank test (ns not significant, ^*^ P<0.05, ^**^ P<0.01).

**Supplemental Fig. 3.**
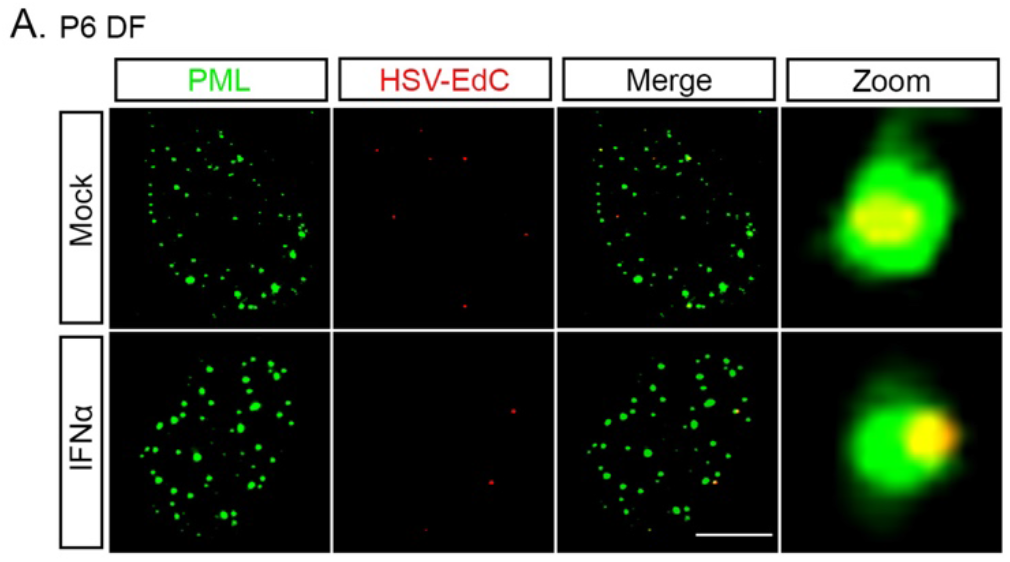
PML-NBs entrap vDNA in the absence of type I IFN during lytic HSV-1 infection of murine dermal fibroblasts. (**A**) Representative images of vDNA foci detected by click chemistry to PML at 60 minutes post-infection in P6 dermal fibroblasts lytically infected with HSV- 1^EdC^ in the presence or absence of IFNα (600 IU/ml).

**Supplemental Fig. 4.**
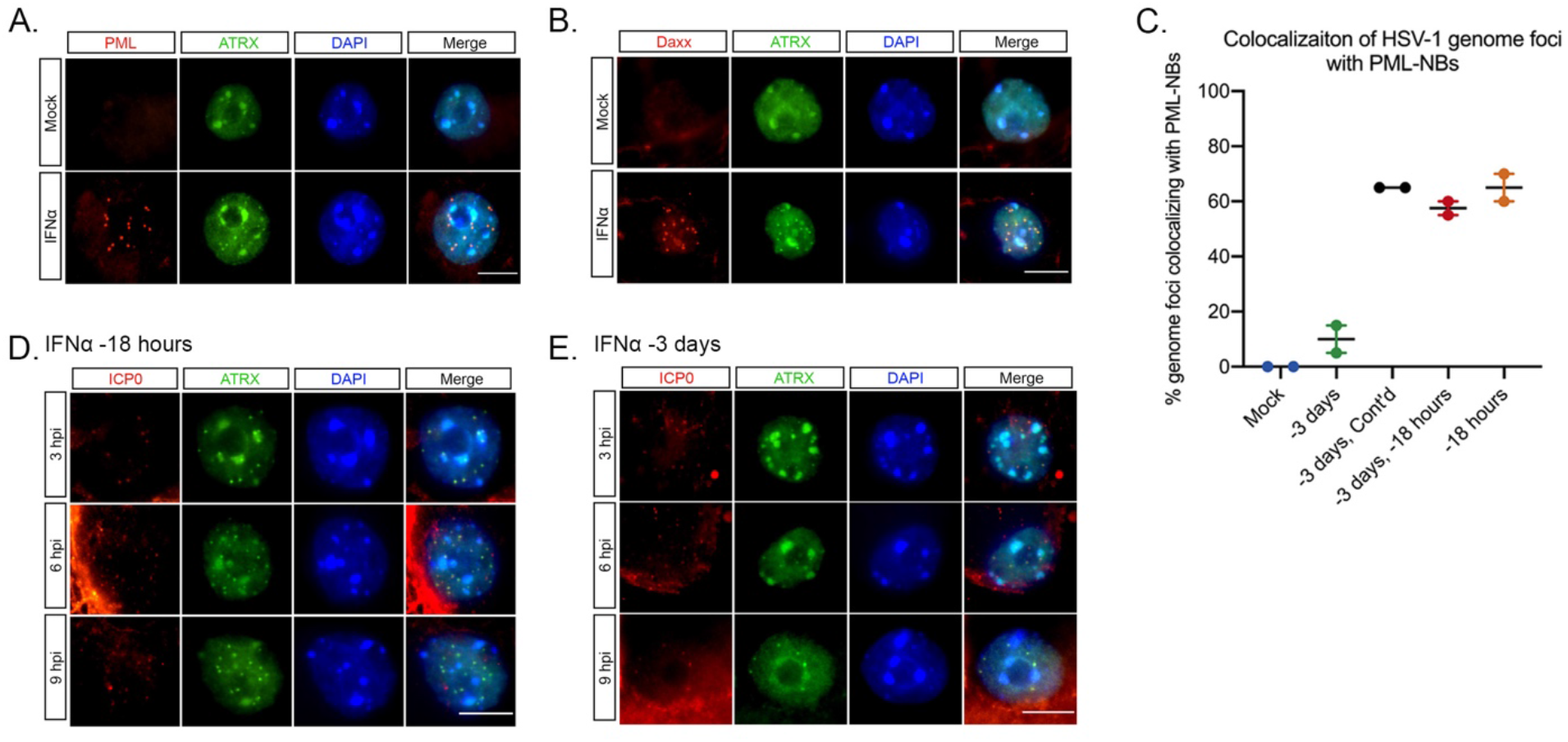
HSV-1 genomes only associate with PML-NBs when type I IFN is present during initial infection. (**A**) Representative images of untreated or IFNα-treated (600 IU/ml) P6 SCG neurons stained for PML and ATRX at 3 days post-treatment. (**B**) Representative images of untreated or IFNα-treated (600 IU/ml) P6 SCG neurons stained for Daxx and ATRX at 3 days post- treatment. (**C**) Percent colocalization of vDNA foci detected by click chemistry to PML at 3 dpi in SCG neurons infected with HSV-1^EdC^ with or without IFNα (600 IU/ml) present at the time of infection. (**D, E**) Representative images of HSV-1-infected P6 SCG neurons treated with IFNα (600 IU/ml) for 18 hours prior to infection or for 18hours at 3 days prior to infection and stained for ICP0 and ATRX at 3, 6 and 9 hours post-infection. Scale bar, 20μm.

## Materials and Methods

### Reagents

Compounds used in the study are as follows: Acycloguanosine, FUDR, LY 294002, Nerve Growth Factor 2.5S (Alomone Labs), Primocin (Invivogen), Aphidicolin (AG Scientific), IFN-α (EMD Millipore IF009), IFN-β (EMD Millipore IF011), IFN-γ (EMD Millipore IF005), IFN-λ2 (PeproTech 250-33); WAY-150138 was kindly provided by Pfizer, Dr. Jay Brown and Dr. Dan Engel at the University of Virginia, and Dr. Lynn Enquist at Princeton University. Compound information and concentrations used can be found below in Table S1.

### Preparation of HSV-1 Virus Stocks

HSV-1 stocks of eGFP-Us11 Patton were grown and titrated on Vero cells obtained from the American Type Culture Collection (Manassas, VA). Cells were maintained in Dulbecco’s Modified Eagle’s Medium (Gibco) supplemented with 10% FetalPlex (Gemini Bio-Products) and 2 mM L-Glutamine. eGFP-Us11 Patton (HSV-1 Patton strain with eGFP reporter protein fused to true late protein Us11 (Benboudjema et al., 2003)) was kindly provided by Dr. Ian Mohr at New York University.

Stayput Us11-GFP was created by inserting an eUs11-GFP tag into the previously created gH-deficient HSV-1 SCgHZ virus (strain SC16) through co-transfection of SCgHZ viral DNA and pSXZY-eGFP-Us11 plasmid (Forrester et al., 1992). Stayput Us11-GFP is propagated and titrated on previously constructed Vero F6 cells, which contain copies of the gH gene under the control of an HSV-1 gD promoter, as described in Forrester et al. (1992). Vero F6s are maintained in Dulbecco’s Modified Eagle’s Medium (Gibco) supplemented with 10% FetaPlex (Gemini BioProducts). They are selected with The supplementation of 250 ug/mL of G418/Geneticin (Gibco).

### Primary Neuronal Cultures

Sympathetic neurons from the Superior Cervical Ganglia (SCG) of post-natal day 0-2 (P0-P2) or adult (P21-P24) CD1 Mice (Charles River Laboratories) were dissected as previously described (Cliffe et al., 2015). Sensory neurons from Trigeminal Ganglia (TG) of post-natal day 0-2 (P0-P2) CD1 mice (Charles River Laboratories) were dissected using the same protocol. Sensory neurons from TG of adult were dissected as previously described (Bertke et al., 2011) with a modified purification protocol using Percoll from the protocol published by Malin et al. (2007). Rodent handling and husbandry were carried out under animal protocols approved by the Animal Care and Use Committee of the University of Virginia (UVA). Ganglia were briefly kept in Leibovitz’s L-15 media with 2.05 mM L-Glutamine before dissociation in Collagenase Type IV (1 mg/mL) followed by Trypsin (2.5 mg/mL) for 20 minutes each at 37 °C. Dissociated ganglia were triturated, and approximately 10,000 neurons per well were plated onto rat tail collagen in a 24-well plate. Sympathetic neurons were maintained in CM1 (Neurobasal® Medium supplemented with PRIME-XV IS21 Neuronal Supplement (Irvine Scientific), 50 ng/mL Mouse NGF 2.5S, 2 mM L-Glutamine, and Primocin). Aphidicolin (3.3 µg/mL) was added to the CM1 for the first five days post-dissection to select against proliferating cells. Sensory neurons were maintained in the same media supplemented with GDNF (50ng/ml; Peprotech 450-44)

### Establishment and Reactivation of Latent HSV-1 Infection in Primary Neurons

Latent HSV-1 infection was established in P6-8 sympathetic neurons from SCGs. Neurons were cultured for at least 24 hours without antimitotic agents prior to infection. The cultures were infected with eGFP-Us11 (Patton recombinant strain of HSV-1 expressing an eGFP reporter fused to true late protein Us11) or StayPut. Neurons were infected at a Multiplicity of Infection (MOI) of 7.5 PFU/cell with eGFP-Us11 and at an MOI of 5 PFU/cell with StayPut (assuming 1.0×10^4^ neurons/well/24-well plate) in DPBS +CaCl_2_ +MgCl_2_ supplemented with 1% Fetal Bovine Serum, 4.5 g/L glucose, and 10 µM Acyclovir (ACV) for 2-3 hours at 37 °C. Post-infection, inoculum was replaced with CM1 containing 50 µM ACV and an anti-mouse IFNAR-1 antibody (Leinco Tech I-1188, 1:1000) for 5-6 days, followed by CM1 without ACV. Reactivation was carried out in DMEM/F12 (Gibco) supplemented with 10% Fetal Bovine Serum, Mouse NGF 2.5S (50 ng/mL) and Primocin. WAY-150138 (10 µg/mL) was added to reactivation cocktail to limit cell-to-cell spread. Reactivation was quantified by counting number of GFP-positive neurons or performing Reverse Transcription Quantitative PCR (RT-qPCR) of HSV-1 lytic mRNAs isolated from the cells in culture.

### Analysis of mRNA expression by reverse-transcription quantitative PCR (RT- qPCR)

To assess relative expression of HSV-1 lytic mRNA, total RNA was extracted from approximately 1.0×10^4^ neurons using the Quick-RNA^™^ Miniprep Kit (Zymo Research) with an on-column DNase I digestion. mRNA was converted to cDNA using the SuperScript IV First-Strand Synthesis system (Invitrogen) using random hexamers for first strand synthesis and equal amounts of RNA (20-30 ng/reaction). To assess viral DNA load, total DNA was extracted from approximately 1.0×10^4^ neurons using the Quick-DNA^™^ Miniprep Plus Kit (Zymo Research). qPCR was carried out using *Power* SYBR^™^ Green PCR Master Mix (Applied Biosystems). The relative mRNA or DNA copy number was determined using the Comparative C_T_ (ΔΔC_T_) method normalized to mRNA or DNA levels in latently infected samples. Viral RNAs were normalized to mouse reference gene GAPDH. All samples were run in duplicate on an Applied Biosystems^™^ QuantStudio^™^ 6 Flex Real-Time PCR System and the mean fold change compared to the reference gene calculated. Primers used are described in Table S2.

### Immunofluorescence

Neurons were fixed for 15 minutes in 4% Formaldehyde and blocked in 5% Bovine Serum Albumin and 0.3% Triton X-100 and incubated overnight in primary antibody. Following primary antibody treatment, neurons were incubated for one hour in Alexa Fluor® 488-, 555-, and 647-conjugated secondary antibodies for multi-color imaging (Invitrogen). Nuclei were stained with Hoechst 33258 (Life Technologies). Images were acquired using an sCMOS charge-coupled device camera (pco.edge) mounted on a Nikon Eclipse Ti Inverted Epifluorescent microscope using NIS-Elements software (Nikon). Images were analyzed and processed using ImageJ.

### Click Chemistry

For EdC-labeled HSV-1 virus infections, an MOI of 5 was used. EdC labelled virus was prepared using a previously described method (McFarlane et al., 2019). Click chemistry was carried out a described previously (Alandijany et al., 2018) with some modifications. Neurons were washed with CSK buffer (10 mM HEPES, 100 mM NaCl, 300 mM Sucrose, 3 mM MgCl_2_, 5 mM EGTA) and simultaneously fixed and permeabilized for 10 minutes in 1.8% methonal-free formaldehyde (0.5% Triton X-100, 1% phenylmethylsulfonyl fluoride (PMSF)) in CSK buffer, then washed twice with PBS before continuing to the click chemistry reaction and immunostaining. Samples were blocked with 3% BSA for 30 minutes, followed by click chemistry using EdC-labelled HSV-1 DNA and the Click-iT EdU Alexa Flour 555 Imaging Kit (ThermoFisher Scientific, C10638) according to the manufacturer’s instructions with AFDye 555 Picolyl Azide (Click Chemistry Tools, 1288). For immunostaining, samples were incubated overnight with primary antibodies in 3% BSA. Following primary antibody treatment, neurons were incubated for one hour in Alexa Fluor® 488- and 647-conjugated secondary antibodies for multi-color imaging (Invitrogen). Nuclei were stained with Hoechst 33258 (Life Technologies). Epifluorescence microscopy images were acquired at 60x using an sCMOS charge-coupled device camera (pco.edge) mounted on a Nikon Eclipse Ti Inverted Epifluorescent microscope using NIS-Elements software (Nikon). Images were analyzed and processed using ImageJ. Confocal microscopy images were acquired using a Zeiss LSM 880 confocal microscope using the 63x Plan-Apochromat oil immersion lens (numerical aperture 1.4) using 405 nm, 488 nm, 543 nm, and 633 nm laser lines. Zen black software (Zeiss) was used for image capture, generating cut mask channels, and calculating weighted colocalization coefficients. Exported images were processed with minimal adjustment using Adobe Photoshop and assembled for presentation using Adobe Illustrator.

### Preparation of Lentiviral Vectors

Lentiviruses expressing shRNA against PML (PML-1 TRCN0000229547, PML-2 TRCN0000229549, PML-3 TRCN0000314605), or a control lentivirus shRNA (Everett et al., 2006) were prepared by co-transfection with psPAX2 and pCMV-VSV-G (Stewart et al., 2003) using the 293LTV packaging cell line (Cell Biolabs). Supernatant was harvested at 40- and 64-hours post-transfection. Sympathetic neurons were transduced overnight in neuronal media containing 8μg/ml protamine and 50μM ACV.

### RNA Sequence Analysis

Reads were checked for quality using FASTQC (v0.11.8), trimmed using BBMAP (v3.8.16b), and aligned to the mouse genome with GENCODE (vM22) annotations using STAR (v2.7.1a). Transcripts per million calculations were performed by RSEM (v1.3.1), the results of which were imported into R (v4.0.2) and Bioconductor (v3.12) using tximport (v1.18.0). Significant genes were called using DESeq2, using fold change cutoffs and pvalue cutoffs of 0.5 and 0.05 respectively. Results were visualized using Heatplus (v2.36.0), PCAtools (v2.2.0), and UpSetR (v1.4.0). Functional enrichment was performed using GSEA and Metascape.

### Statistical Analysis

Power analysis was used to determine the appropriate sample sizes for statistical analysis. All statistical analysis was performed using Prism V8.4. An unpaired t-test was used for all experiments where the group size was 2. All other experiments were analyzed using a one-way ANOVA with a Tukey’s multiple comparison. Specific analyses are included in the figure legends. For all reactivation experiments measuring GFP expression, viral DNA, gene expression or DNA load, individual biological replicates were plotted (an individual well of primary neurons) and all experiments were repeated from pools of neurons from at least 3 litters.

## Supplemental Materials and Methods Tables

**Table S1:**
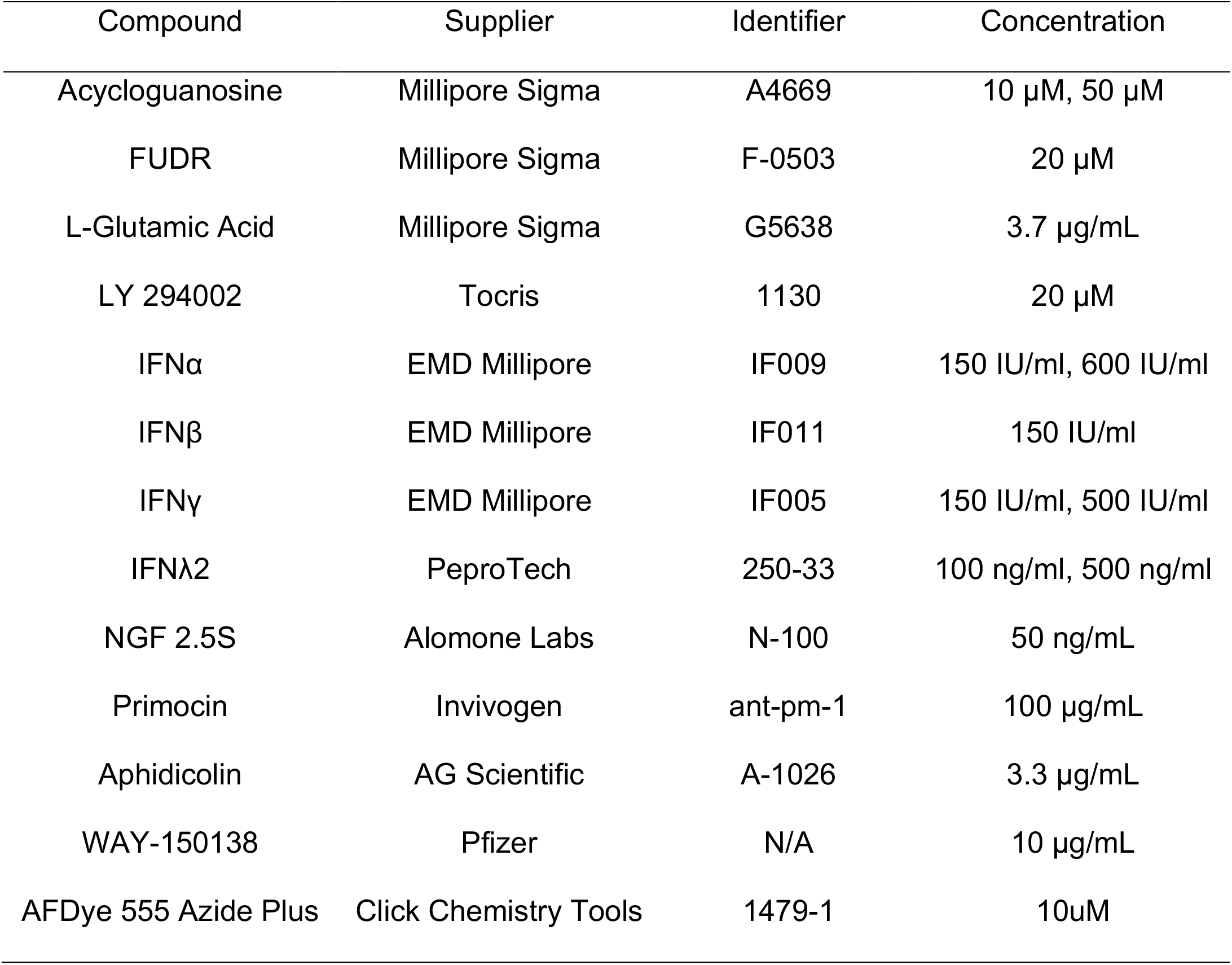
Compounds Used and Concentrations

| Compound | Supplier | Identifier | Concentration |
| --- | --- | --- | --- |
| Acycloguanosine | Millipore Sigma | A4669 | 10 µM, 50 µM |
| FUDR | Millipore Sigma | F-0503 | 20 µM |
| L-Glutamic Acid | Millipore Sigma | G5638 | 3.7 µg/mL |
| LY 294002 | Tocris | 1130 | 20 µM |
| IFNα | EMD Millipore | IF009 | 150 IU/ml, 600 IU/ml |
| IFNβ | EMD Millipore | IF011 | 150 IU/ml |
| IFNγ | EMD Millipore | IF005 | 150 IU/ml, 500 IU/ml |
| IFNλ2 | PeproTech | 250-33 | 100 ng/ml, 500 ng/ml |
| NGF 2.5S | Alomone Labs | N-100 | 50 ng/mL |
| Primocin | Invivogen | ant-pm-1 | 100 µg/mL |
| Aphidicolin | AG Scientific | A-1026 | 3.3 µg/mL |
| WAY-150138 | Pfizer | N/A | 10 µg/mL |
| AFDye 555 Azide Plus | Click Chemistry Tools | 1479-1 | 10uM |

**Table S2:**
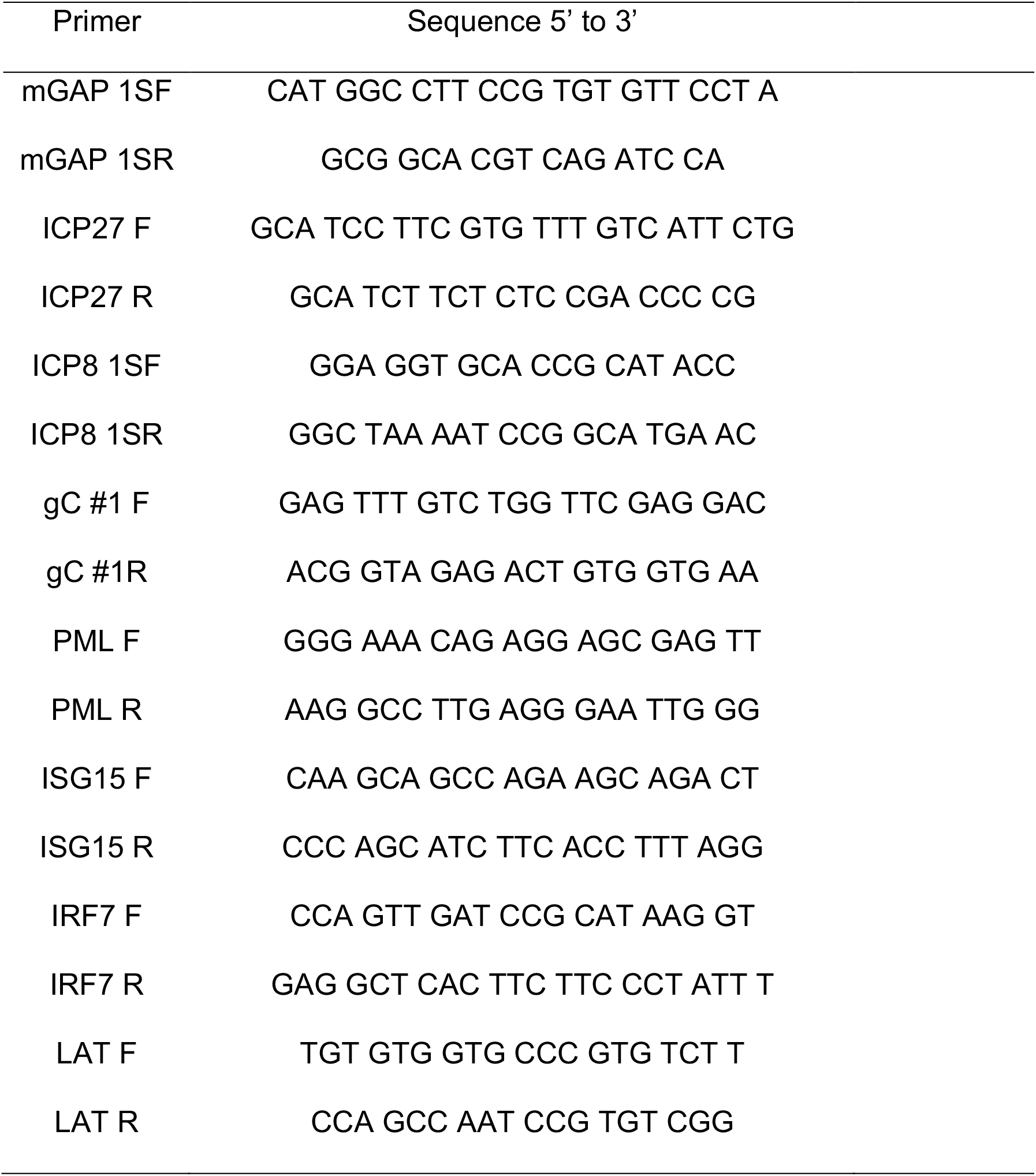
Primers Used for RT-qPCR

**Table S4:** Antibodies Used for Immunofluorescence and Concentrations

| Antibody | Supplier | Identifier | Concentration |
| --- | --- | --- | --- |
| Ms PML | EMD Millipore | MAB3738 | 1:200 |
| Ch Beta-III Tubulin | Millipore sigma | AB9354 | 1:500 |
| Rb ATRX (H-300) | Santa Cruz Bio | sc-15408 | 1:250 |
| Ms Daxx (H-7) | Santa Cruz Bio | sc-8043 | 1:250 |
| Rb STAT1 (D1K9Y) | CST | mAb 14994 | 1:400 |
| Ms Mx1/2/3 | Santa Cruz Bio | sc-166412 | 1:250 |
| Ms HSV-1 ICP0 | East Coast Bio | H1A027 | 1:200 |
| F(ab') Goat anti Mouse IgG (H+L) Alexa Fluor® 555 | Thermo Fisher | A21425 | 1:1000 |
| F(ab')2 Goat anti-Rabbit IgG (H+L) Alexa Fluor® 488 | Thermo Fisher | A11070 | 1:1000 |
| F(ab')2 Goat anti-Rabbit IgG (H+L) Alexa Fluor® 488 | Thermo Fisher | A11017 | 1:1000 |
| Goat anti Chicken IgY (H+L) Alexa Fluor® 647 | abcam | ab150175 | 1:1000 |

## References

Alandijany, T., Roberts, A. P. E., Conn, K. L., Loney, C., Mcfarlane, S., Orr, A. & Boutell, C. 2018. Distinct temporal roles for the promyelocytic leukaemia (PML) protein in the sequential regulation of intracellular host immunity to HSV-1 infection. PLoS Pathogens, 14, e1006769–36.

Barragan-Iglesias, P., Franco-Enzastiga, U., Jeevakumar, V., Shiers, S., Wangzhou, A., Granados-Soto, V., Campbell, Z. T., Dussor, G. & Price, T. J. 2020. Type I Interferons Act Directly on Nociceptors to Produce Pain Sensitization: Implications for Viral Infection-Induced Pain. J Neurosci, 40, 3517–3532.

Benboudjema, L., Mulvey, M., Gao, Y., Pimplikar, S. W. & Mohr, I. 2003. Association of the herpes simplex virus type 1 Us11 gene product with the cellular kinesin light-chain-related protein PAT1 results in the redistribution of both polypeptides. J Virol, 77, 9192–203.

Bernardi, R. & Pandolfi, P. P. 2007. Structure, dynamics and functions of promyelocytic leukaemia nuclear bodies. Nat Rev Mol Cell Biol, 8, 1006–16.

Bertke, A. S., Swanson, S. M., Chen, J., Imai, Y., Kinchington, P. R. & Margolis, T. P. 2011. A5-positive primary sensory neurons are nonpermissive for productive infection with herpes simplex virus 1 in vitro. J Virol, 85, 6669–77.

Bishop, C. L., Ramalho, M., Nadkarni, N., May Kong, W., Higgins, C. F. & Krauzewicz, N. 2006. Role for centromeric heterochromatin and PML nuclear bodies in the cellular response to foreign DNA. Mol Cell Biol, 26, 2583–94.

Bloom, D. C. 2016. Alphaherpesvirus Latency: A Dynamic State of Transcription and Reactivation. Adv Virus Res, 94, 53–80.

Boutell, C., Cuchet-Lourenco, D., Vanni, E., Orr, A., Glass, M., Mcfarlane, S. & Everett, R. D. 2011. A viral ubiquitin ligase has substrate preferential SUMO targeted ubiquitin ligase activity that counteracts intrinsic antiviral defence. PLoS Pathog, 7, e1002245.

Boutell, C., Sadis, S. & Everett, R. D. 2002. Herpes simplex virus type 1 immediate-early protein ICP0 and is isolated RING finger domain act as ubiquitin E3 ligases in vitro. J Virol, 76, 841–50.

Branco, F. J. & Fraser, N. W. 2005. Herpes simplex virus type 1 latency-associated transcript expression protects trigeminal ganglion neurons from apoptosis. J Virol, 79, 9019–25.

Bucevicius, J., Keller-Findeisen, J., Gilat, T., Hell, S. W. & Lukinavicius, G. 2019. Rhodamine-Hoechst positional isomers for highly efficient staining of heterochromatin. Chem Sci, 10, 1962–1970.

Cabral, J. M., Oh, H. S. & Knipe, D. M. 2018. ATRX promotes maintenance of herpes simplex virus heterochromatin during chromatin stress. Elife, 7.

Camarena, V. 2011. Nerve Growth Factor Signaling maintain HSV latency. 1–254.

Carr, D. J., Veress, L. A., Noisakran, S. & Campbell, I. L. 1998. Astrocyte-targeted expression of IFN-alpha1 protects mice from acute ocular herpes simplex virus type 1 infection. J Immunol, 161, 4859–65.

Catez, F., Picard, C., Held, K., Gross, S., Rousseau, A., Theil, D., Sawtell, N., Labetoulle, M. & Lomonte, P. 2012. HSV-1 genome subnuclear positioning and associations with host-cell PML-NBs and centromeres regulate LAT locus transcription during latency in neurons. PLoS Pathog, 8, e1002852.

Chang, J. Y., Martin, D. P. & Johnson, E. M., Jr. 1990. Interferon suppresses sympathetic neuronal cell death caused by nerve growth factor deprivation. J Neurochem, 55, 436–45.

Chelbi-Alix, M. K. & De The, H. 1999. Herpes virus induced proteasome-dependent degradation of the nuclear bodies-associated PML and Sp100 proteins. Oncogene, 18, 935–41.

Chelbi-Alix, M. K., Pelicano, L., Quignon, F., Koken, M. H., Venturini, L., Stadler, M., Pavlovic, J., Degos, L. & De The, H. 1995. Induction of the PML protein by interferons in normal and APL cells. Leukemia, 9, 2027–33.

Chen, S. H., Kramer, M. F., Schaffer, P. A. & Coen, D. M. 1997. A viral function represses accumulation of transcripts from productive-cycle genes in mouse ganglia latently infected with herpes simplex virus. Journal of Virology, 71, 5878–5884.

Chen, Y., Wright, J., Meng, X. & Leppard, K. N. 2015. Promyelocytic Leukemia Protein Isoform II Promotes Transcription Factor Recruitment To Activate Interferon Beta and Interferon-Responsive Gene Expression. Mol Cell Biol, 35, 1660–72.

Cliffe, A. R., Arbuckle, J. H., Vogel, J. L., Geden, M. J., Rothbart, S. B., Cusack, C. L., Strahl, B. D., Kristie, T. M. & Deshmukh, M. 2015. Neuronal Stress Pathway Mediating a Histone Methyl/Phospho Switch Is Required for Herpes Simplex Virus Reactivation. Cell Host Microbe, 18, 649–58.

Cliffe, A. R., Coen, D. M. & Knipe, D. M. 2013. Kinetics of facultative heterochromatin and polycomb group protein association with the herpes simplex viral genome during establishment of latent infection. MBio, 4, e00590-12-e00590-12.

Cliffe, A. R., Garber, D. A. & Knipe, D. M. 2009. Transcription of the herpes simplex virus latency-associated transcript promotes the formation of facultative heterochromatin on lytic promoters. J Virol, 83, 8182–90.

Cliffe, A. R. & Wilson, A. C. 2017. Restarting Lytic Gene Transcription at the Onset of Herpes Simplex Virus Reactivation. J Virol, 91, e01419-16-6.

Clynes, D., Higgs, D. R. & Gibbons, R. J. 2013. The chromatin remodeller ATRX: a repeat offender in human disease. Trends in Biochemical Sciences, 38, 461–466.

Cohen, C., Corpet, A., Roubille, S., Maroui, M. A., Poccardi, N., Rousseau, A., Kleijwegt, C., Binda, O., Texier, P., Sawtell, N., Labetoulle, M. & Lomonte, P. 2018. Promyelocytic leukemia (PML) nuclear bodies (NBs) induce latent/quiescent HSV-1 genomes chromatinization through a PML NB/Histone H3.3/H3.3 Chaperone Axis. PLoS Pathog, 14, e1007313.

Croxton, R., Puto, L. A., De Belle, I., Thomas, M., Torii, S., Hanaii, F., Cuddy, M. & Reed, J. C. 2006. Daxx represses expression of a subset of antiapoptotic genes regulated by nuclear factor-kappaB. Cancer Res, 66, 9026–35.

Cuchet-Lourenco, D., Vanni, E., Glass, M., Orr, A. & Everett, R. D. 2012. Herpes simplex virus 1 ubiquitin ligase ICP0 interacts with PML isoform I and induces its SUMO-independent degradation. J Virol, 86, 11209–22.

Cuddy, S. R., Schinlever, A. R., Dochnal, S., Seegren, P. V., Suzich, J., Kundu, P., Downs, T. K., Farah, M., Desai, B. N., Boutell, C. & Cliffe, A. R. 2020. Neuronal hyperexcitability is a DLK-dependent trigger of herpes simplex virus reactivation that can be induced by IL-1. Elife, 9.

Du, T., Zhou, G. & Roizman, B. 2011. HSV-1 gene expression from reactivated ganglia is disordered and concurrent with suppression of latency-associated transcript and miRNAs. Proceedings of the National Academy of Sciences, 108, 18820–18824.

Efstathiou, S. & Preston, C. M. 2005. Towards an understanding of the molecular basis of herpes simplex virus latency. Virus Res, 111, 108–19.

Everett, R. D. & Chelbi-Alix, M. K. 2007. PML and PML nuclear bodies: implications in antiviral defence. Biochimie, 89, 819–30.

Everett, R. D., Freemont, P., Saitoh, H., Dasso, M., Orr, A., Kathoria, M. & Parkinson, J. 1998. The disruption of ND10 during herpes simplex virus infection correlates with the Vmw110- and proteasome-dependent loss of several PML isoforms. J Virol, 72, 6581–91.

Everett, R. D. & Maul, G. G. 1994. HSV-1 IE protein Vmw110 causes redistribution of PML. EMBO J, 13, 5062–9.

Everett, R. D., Rechter, S., Papior, P., Tavalai, N., Stamminger, T. & Orr, A. 2006. PML contributes to a cellular mechanism of repression of herpes simplex virus type 1 infection that is inactivated by ICP0. J Virol, 80, 7995–8005.

Forrester, A., Farrell, H., Wilkinson, G., Kaye, J., Davis-Poynter, N. & Minson, T. 1992. Construction and properties of a mutant of herpes simplex virus type 1 with glycoprotein H coding sequences deleted. Journal of Virology, 66, 341–348.

Garrick, D., Samara, V., Mcdowell, T. L., Smith, A. J., Dobbie, L., Higgs, D. R. & Gibbons, R. J. 2004. A conserved truncated isoform of the ATR-X syndrome protein lacking the SWI/SNF-homology domain. Gene, 326, 23–34.

Gordon, Y. J., Romanowski, E. G., Araullo-Cruz, T. & Kinchington, P. R. 1995. The proportion of trigeminal ganglionic neurons expressing herpes simplex virus type 1 latency-associated transcripts correlates to reactivation in the New Zealand rabbit ocular model. Graefes Arch Clin Exp Ophthalmol, 233, 649–54.

Gray, P. A., Fu, H., Luo, P., Zhao, Q., Yu, J., Ferrari, A., Tenzen, T., Yuk, D. I., Tsung, E. F., Cai, Z., Alberta, J. A., Cheng, L. P., Liu, Y., Stenman, J. M., Valerius, M. T., Billings, N., Kim, H. A., Greenberg, M. E., Mcmahon, A. P., Rowitch, D. H., Stiles, C. D. & Ma, Q. 2004. Mouse brain organization revealed through direct genome-scale TF expression analysis. Science, 306, 2255–7.

Greger, J. G., Katz, R. A., Ishov, A. M., Maul, G. G. & Skalka, A. M. 2005. The cellular protein daxx interacts with avian sarcoma virus integrase and viral DNA to repress viral transcription. J Virol, 79, 4610–8.

Grotzinger, T., Jensen, K. & Will, H. 1996. The interferon (IFN)-stimulated gene Sp100 promoter contains an IFN-gamma activation site and an imperfect IFN-stimulated response element which mediate type I IFN inducibility. J Biol Chem, 271, 25253–60.

Guo, A., Salomoni, P., Luo, J., Shih, A., Zhong, S., Gu, W. & Pandolfi, P. P. 2000. The function of PML in p53-dependent apoptosis. Nat Cell Biol, 2, 730–6.

Hall, M. H., Magalska, A., Malinowska, M., Ruszczycki, B., Czaban, I., Patel, S., Ambrozek-Latecka, M., Zolocinska, E., Broszkiewicz, H., Parobczak, K., Nair, R. R., Rylski, M., Pawlak, R., Bramham, C. R. & Wilczynski, G. M. 2016. Localization and regulation of PML bodies in the adult mouse brain. Brain Struct Funct, 221, 2511–25.

Hendricks, R. L., Weber, P. C., Taylor, J. L., Koumbis, A., Tumpey, T. M. & Glorioso, J. C. 1991. Endogenously produced interferon alpha protects mice from herpes simplex virus type 1 corneal disease. J Gen Virol, 72 (Pt 7), 1601–10.

Hochrein, H., Schlatter, B., O’Keeffe, M., Wagner, C., Schmitz, F., Schiemann, M., Bauer, S., Suter, M. & Wagner, H. 2004. Herpes simplex virus type-1 induces IFN-alpha production via Toll-like receptor 9-dependent and - independent pathways. Proc Natl Acad Sci U S A, 101, 11416–21.

Ives, A. M. & Bertke, A. S. 2017. Stress Hormones Epinephrine and Corticosterone Selectively Modulate Herpes Simplex Virus 1 (HSV-1) and HSV-2 Productive Infections in Adult Sympathetic, but Not Sensory, Neurons. Journal of Virology, 91, e00582-17-12.

Jones, C. A., Fernandez, M., Herc, K., Bosnjak, L., Miranda-Saksena, M.,Boadle, R. A. & Cunningham, A. 2003. Herpes simplex virus type 2 induces rapid cell death and functional impairment of murine dendritic cells in vitro. J Virol, 77, 11139–49.

Kamada, R., Yang, W., Zhang, Y., Patel, M. C., Yang, Y., Ouda, R., Dey, A., Wakabayashi, Y., Sakaguchi, K., Fujita, T., Tamura, T., Zhu, J. & Ozato, K. 2018. Interferon stimulation creates chromatin marks and establishes transcriptional memory. Proceedings of the National Academy of Sciences, 115, E9162– E9171.

Katzenell, S. & Leib, D. A. 2016. Herpes Simplex Virus and Interferon Signaling Induce Novel Autophagic Clusters in Sensory Neurons. J Virol, 90, 4706–4719.

Kim, J. Y., Mandarino, A., Chao, M. V., Mohr, I. & Wilson, A. C. 2012. Transient reversal of episome silencing precedes VP16-dependent transcription during reactivation of latent HSV-1 in neurons. PLoS Pathog, 8, e1002540.

Kim, Y. E. & Ahn, J. H. 2015. Positive role of promyelocytic leukemia protein in type I interferon response and its regulation by human cytomegalovirus. PLoS Pathog, 11, e1004785.

Knipe, D. M. & Cliffe, A. 2008. Chromatin control of herpes simplex virus lytic and latent infection. Nat Rev Microbiol, 6, 211–21.

Kobayashi, M., Wilson, A. C., Chao, M. V. & Mohr, I. 2012. Control of viral latency in neurons by axonal mTOR signaling and the 4E-BP translation repressor. Genes Dev, 26, 1527–32.

Kramer, M. F. & Coen, D. M. 1995. Quantification of transcripts from the ICP4 and thymidine kinase genes in mouse ganglia latently infected with herpes simplex virus. J Virol, 69, 1389–99.

Kreit, M., Paul, S., Knoops, L., De Cock, A., Sorgeloos, F. & Michiels, T. 2014. Inefficient type I interferon-mediated antiviral protection of primary mouse neurons is associated with the lack of apolipoprotein l9 expression. J Virol, 88, 3874–84.

Kwiatkowski, D. L., Thompson, H. W. & Bloom, D. C. 2009. The polycomb group protein Bmi1 binds to the herpes simplex virus 1 latent genome and maintains repressive histone marks during latency. J Virol, 83, 8173–81.

Lallemand-Breitenbach, V. & De The, H. 2010. PML nuclear bodies. Cold Spring Harb Perspect Biol, 2, a000661.

Lewis, P. W., Elsaesser, S. J., Noh, K.-M., Stadler, S. C. & Allis, C. D. 2010. Daxx is an H3.3-specific histone chaperone and cooperates with ATRX in replication- independent chromatin assembly at telomeres. Proceedings of the National Academy of Sciences of the United States of America, 107, 14075–14080.

Li, H., Zhang, J., Kumar, A., Zheng, M., Atherton, S. S. & Yu, F. S. 2006. Herpes simplex virus 1 infection induces the expression of proinflammatory cytokines, interferons and TLR7 in human corneal epithelial cells. Immunology, 117, 167–76.

Liang, Y., Vogel, J. L., Arbuckle, J. H., Rai, G., Jadhav, A., Simeonov, A., Maloney, D. J. & Kristie, T. M. 2013. Targeting the JMJD2 histone demethylases to epigenetically control herpesvirus infection and reactivation from latency. Sci Transl Med, 5, 167ra5.

Liang, Y., Vogel, J. L., Narayanan, A., Peng, H. & Kristie, T. M. 2009. Inhibition of the histone demethylase LSD1 blocks alpha-herpesvirus lytic replication and reactivation from latency. Nat Med, 15, 1312–7.

Lukashchuk, V. & Everett, R. D. 2010. Regulation of ICP0-null mutant herpes simplex virus type 1 infection by ND10 components ATRX and hDaxx. J Virol, 84, 4026–40.

Ma, J. Z., Russell, T. A., Spelman, T., Carbone, F. R. & Tscharke, D. C. 2014. Lytic Gene Expression Is Frequent in HSV-1 Latent Infection and Correlates with the Engagement of a Cell-Intrinsic Transcriptional Response. PLoS Pathogens, 10, e1004237.

Malin, S. A., Davis, B. M. & Molliver, D. C. 2007. Production of dissociated sensory neuron cultures and considerations for their use in studying neuronal function and plasticity. Nat Protoc, 2, 152–60.

Maroui, M. A., CallÉ, A., Cohen, C., Streichenberger, N., Texier, P., Takissian, J., Rousseau, A., Poccardi, N., Welsch, J., Corpet, A., Schaeffer, L., Labetoulle, M. & Lomonte, P. 2016. Latency Entry of Herpes Simplex Virus 1 Is Determined by the Interaction of Its Genome with the Nuclear Environment. PLoS Pathogens, 12, e1005834–28.

Maul, G. G. 1998. Nuclear domain 10, the site of DNA virus transcription and replication. Bioessays, 20, 660–7.

Maul, G. G., Ishov, A. M. & Everett, R. D. 1996. Nuclear domain 10 as preexisting potential replication start sites of herpes simplex virus type-1. Virology, 217, 67–75.

Mcfarlane, S., Orr, A., Roberts, A. P. E., Conn, K. L., Iliev, V., Loney, C., DA SILVA Filipe, A., Smollett, K., Gu, Q., Robertson, N., Adams, P. D., Rai, T. S. & Boutell, C. 2019. The histone chaperone HIRA promotes the induction of host innate immune defences in response to HSV-1 infection. PLOS Pathogens, 15, e1007667.

Mikloska, Z. & Cunningham, A. L. 2001. Alpha and gamma interferons inhibit herpes simplex virus type 1 infection and spread in epidermal cells after axonal transmission. J Virol, 75, 11821–6.

Mikloska, Z., Danis, V. A., Adams, S., Lloyd, A. R., Adrian, D. L. & Cunningham, A. L. 1998. In vivo production of cytokines and beta (C-C) chemokines in human recurrent herpes simplex lesions--do herpes simplex virus-infected keratinocytes contribute to their production? J Infect Dis, 177, 827–38.

Moorlag, S., Roring, R. J., Joosten, L. A. B. & Netea, M. G. 2018. The role of the interleukin-1 family in trained immunity. Immunol Rev, 281, 28–39.

Muller, S., Matunis, M. J. & Dejean, A. 1998. Conjugation with the ubiquitin-related modifier SUMO-1 regulates the partitioning of PML within the nucleus. EMBO J, 17, 61– 70.

Nicoll, M. P., Hann, W., Shivkumar, M., Harman, L. E., Connor, V., Coleman, H. M., Proenca, J. T. & Efstathiou, S. 2016. The HSV-1 Latency-Associated Transcript Functions to Repress Latent Phase Lytic Gene Expression and Suppress Virus Reactivation from Latently Infected Neurons. PLoS Pathog, 12, e1005539.

Noh, K. M., Maze, I., Zhao, D., Xiang, B., Wenderski, W., Lewis, P. W., Shen, L., Li, H. & Allis, C. D. 2015. ATRX tolerates activity-dependent histone H3 methyl/phos switching to maintain repetitive element silencing in neurons. Proc Natl Acad Sci U S A, 112, 6820–7.

Proenca, J. T., Coleman, H. M., Connor, V., Winton, D. J. & Efstathiou, S. 2008. A historical analysis of herpes simplex virus promoter activation in vivo reveals distinct populations of latently infected neurones. Journal of General Virology, 89, 2965–2974.

Quignon, F., De Bels, F., Koken, M., Feunteun, J., Ameisen, J. C. & De The, H. 1998. PML induces a novel caspase-independent death process. Nat Genet, 20, 259–65.

Rasmussen, S. B., Jensen, S. B., Nielsen, C., Quartin, E., Kato, H., Chen, Z. J., Silverman, R. H., Akira, S. & Paludan, S. R. 2009. Herpes simplex virus infection is sensed by both Toll-like receptors and retinoic acid-inducible gene- like receptors, which synergize to induce type I interferon production. J Gen Virol, 90, 74–8.

Rasmussen, S. B., Sorensen, L. N., Malmgaard, L., Ank, N., Baines, J. D., Chen, Z. J. & Paludan, S. R. 2007. Type I interferon production during herpes simplex virus infection is controlled by cell-type-specific viral recognition through Toll-like receptor 9, the mitochondrial antiviral signaling protein pathway, and novel recognition systems. J Virol, 81, 13315–24.

Regad, T., Bellodi, C., Nicotera, P. & Salomoni, P. 2009. The tumor suppressor Pml regulates cell fate in the developing neocortex. Nat Neurosci, 12, 132–40.

Rosato, P. C. & Leib, D. A. 2014. Intrinsic innate immunity fails to control herpes simplex virus and vesicular stomatitis virus replication in sensory neurons and fibroblasts. J Virol, 88, 9991–10001.

Sainz, B., Jr. & Halford, W. P. 2002. Alpha/Beta interferon and gamma interferon synergize to inhibit the replication of herpes simplex virus type 1. J Virol, 76, 11541–50.

Sawtell, N. M. 1997. Comprehensive quantification of herpes simplex virus latency at the single-cell level. J Virol, 71, 5423–31.

Scherer, M., Otto, V., Stump, J. D., Klingl, S., Muller, R., Reuter, N., Muller, Y. A., Sticht, H. & Stamminger, T. 2016. Characterization of Recombinant Human Cytomegaloviruses Encoding IE1 Mutants L174P and 1-382 Reveals that Viral Targeting of PML Bodies Perturbs both Intrinsic and Innate Immune Responses. J Virol, 90, 1190–205.

Shalginskikh, N., Poleshko, A., Skalka, A. M. & Katz, R. A. 2013. Retroviral DNA methylation and epigenetic repression are mediated by the antiviral host protein Daxx. J Virol, 87, 2137–50.

Song, R., Koyuncu, O. O., Greco, T. M., Diner, B. A., Cristea, I. M. & Enquist, L. W. 2016. Two Modes of the Axonal Interferon Response Limit Alphaherpesvirus Neuroinvasion. mBio, 7, e02145–15.

Stadler, M., Chelbi-Alix, M. K., Koken, M. H., Venturini, L., Lee, C., Saib, A., Quignon, F., Pelicano, L., Guillemin, M. C., Schindler, C. & ET AL. 1995. Transcriptional induction of the PML growth suppressor gene by interferons is mediated through an ISRE and a GAS element. Oncogene, 11, 2565–73.

Stevens, J. G., Wagner, E. K., Devi-Rao, G. B., Cook, M. L. & Feldman, L. T. 1987. RNA complementary to a herpesvirus alpha gene mRNA is prominent in latently infected neurons. Science, 235, 1056–1059.

Stewart, S. A., Dykxhoorn, D. M., Palliser, D., Mizuno, H., Yu, E. Y., An, D. S., Sabatini, D. M., Chen, I. S., Hahn, W. C., Sharp, P. A., Weinberg, R. A. & Novina, C. D. 2003. Lentivirus-delivered stable gene silencing by RNAi in primary cells. Rna, 9, 493–501.

Suzich, J. B. & Cliffe, A. R. 2018. Strength in diversity: Understanding the pathways to herpes simplex virus reactivation. Virology, 522, 81–91.

Thompson, R. L. & Sawtell, N. M. 1997. The herpes simplex virus type 1 latency-associated transcript gene regulates the establishment of latency. J Virol, 71, 5432–40.

Thompson, R. L. & Sawtell, N. M. 2001. Herpes simplex virus type 1 latency-associated transcript gene promotes neuronal survival. J Virol, 75, 6660–75.

Ulbricht, T., Alzrigat, M., Horch, A., Reuter, N., Von Mikecz, A., Steimle, V., Schmitt, E., Kramer, O. H., Stamminger, T. & Hemmerich, P. 2012. PML promotes MHC class II gene expression by stabilizing the class II transactivator. J Cell Biol, 199, 49–63.

Van Zeijl, M., Fairhurst, J., Jones, T. R., Vernon, S. K., Morin, J., Larocque, J., Feld, B., O’Hara, B., Bloom, J. D. & Johann, S. V. 2000. Novel class of thiourea compounds that inhibit herpes simplex virus type 1 DNA cleavage and encapsidation: resistance maps to the UL6 gene. J Virol, 74, 9054–61.

Villagra, N. T., Berciano, J., Altable, M., Navascues, J., Casafont, I., Lafarga, M. & Berciano, M. T. 2004. PML bodies in reactive sensory ganglion neurons of the Guillain-Barre syndrome. Neurobiol Dis, 16, 158–68.

Wang, J., Shiels, C., Sasieni, P., Wu, P. J., Islam, S. A., Freemont, P. S. & Sheer, D. 2004. Promyelocytic leukemia nuclear bodies associate with transcriptionally active genomic regions. J Cell Biol, 164, 515–26.

Wang, Q.-Y., Zhou, C., Johnson, K. E., Colgrove, R. C., Coen, D. M. & Knipe, D. M. 2005. Herpesviral latency-associated transcript gene promotes assembly of heterochromatin on viral lytic-gene promoters in latent infection. Proceedings of the National Academy of Sciences, 102, 16055–16059.

Wang, Z. G., Ruggero, D., Ronchetti, S., Zhong, S., Gaboli, M., Rivi, R. & Pandolfi, P. P. 1998. PML is essential for multiple apoptotic pathways. Nat Genet, 20, 266–72.

Wilcox, C. L. & Johnson, E. M. 1987. Nerve growth factor deprivation results in the reactivation of latent herpes simplex virus in vitro. Journal of Virology, 61, 2311–2315.

Wilcox, C. L., Smith, R. L., Freed, C. R. & Johnson, E. M. 1990. Nerve growth factor- dependence of herpes simplex virus latency in peripheral sympathetic and sensory neurons in vitro. Journal of Neuroscience, 10, 1268–1275.

Woulfe, J., Gray, D., Prichett-Pejic, W., Munoz, D. G. & Chretien, M. 2004. Intranuclear rodlets in the substantia nigra: interactions with marinesco bodies, ubiquitin, and promyelocytic leukemia protein. J Neuropathol Exp Neurol, 63, 1200–7.

Xu, P. & Roizman, B. 2017. The SP100 component of ND10 enhances accumulation of PML and suppresses replication and the assembly of HSV replication compartments. Proc Natl Acad Sci U S A, 114, E3823–E3829.

Yordy, B., Iijima, N., Huttner, A., Leib, D. & Iwasaki, A. 2012. A neuron-specific role for autophagy in antiviral defense against herpes simplex virus. Cell Host and Microbe, 12, 334–345.

Zhong, S., Salomoni, P. & Pandolfi, P. P. 2000. The transcriptional role of PML and the nuclear body. Nat Cell Biol, 2, E85–90.

